# Novel strand exchange activity of the human PALB2 DNA Binding Domain and its critical role for DNA repair in cells

**DOI:** 10.1101/495192

**Authors:** Jaigeeth Deveryshetty, Mikhail Ryzhikov, Nadine Brahiti, Thibaut Peterlini, Graham Dellaire, Jean-Yves Masson, Sergey Korolev

**Affiliations:** Edward A. Doisy Department of Biochemistry and Molecular Biology, Saint Louis University School of Medicine, St. Louis, MO 63104, USA; Genome Stability Laboratory, CHU de Québec Research Center, HDQ Pavilion, Oncology Axis, 9 McMahon, Québec City, QC G1R 2J6, Canada; Department of Pathology, Dalhousie University, Halifax Nova Scotia, Canada

## Abstract

Breast cancer associated proteins 1 and 2 (BRCA1, −2) and partner and localizer of BRCA2 (PALB2) protein are tumor suppressors linked to a spectrum of malignancies, including breast cancer and Fanconi anemia. They stimulate RAD51 recombinase during homology-directed repair (HDR). Along with being a hub for a protein interaction network, PALB2 interacts with DNA. The mechanism of PALB2 DNA binding and its function are poorly understood. We identified a major DNA-binding site in PALB2, mutation of which reduces the RAD51 foci formation and the overall HDR efficiency in cells by 50%. PALB2 N-terminal DNA-binding domain (N-DBD) stimulates the RAD51 strand exchange reaction. Surprisingly, it promotes the strand exchange without RAD51. Moreover, N-DBD stimulates the inverse strand exchange and can use both DNA and RNA substrates. Our data reveal a versatile DNA interaction property of PALB2 and demonstrate a critical role of PALB2 DNA binding for chromosome repair in cells.

## INTRODUCTION

Breast cancer associated proteins 1 and 2 (BRCA1, −2) regulate an efficient non-mutagenic pathway of chromosome break repair and are described as guardians of chromosomal integrity (Venkitaraman, 2014). They initiate RAD51-mediated homologous recombination (HR) (Davies et al., 2001; Moynahan et al., 2001; Sharan et al., 1997; Venkitaraman, 2000) and facilitate restart of stalled replication (Badie et al., 2010; Lomonosov et al., 2003; Schlacher et al., 2011). The partner and localizer of BRCA2 (PALB2) protein was discovered as a protein forming a complex with BRCA2 and regulating BRCA2 activity (Xia et al., 2006). Similarly, to BRCA proteins, PALB2 is an essential mammalian protein linked to a similar spectrum of cancers and Fanconi anemia (Pauty et al., 2014; Xia et al., 2007). PALB2 C-terminal WD40 domain interacts with BRCA2 (Oliver et al., 2009; Xia et al., 2006) while the N-terminus forms a complex with BRCA1 (Zhang et al., 2009a; Zhang et al., 2009b). The latter localizes at double stranded DNA break (DSB) sites at earlier stage of repair, inhibiting an alternative pathway of non-homologous end joining and initiating homology-directed repair (HDR) through interactions with PALB2/BRCA2/RAD51(Prakash et al., 2015).

PALB2 is often described as the hub for a network of tumor suppressors involved in DNA repair (Park et al., 2014b; Sy et al., 2009b). In addition to BRCA1 and −2 interactions, it contains a chromatin-association motif (ChAM) in its central region, responsible for PALB2 association with chromatin through the nucleosome core histones H3 and H2B (Bleuyard et al., 2012). It interacts with MRG15 protein, a component of histone acetyltransferase-deacetylase complexes (Hayakawa et al., 2010; Sy et al., 2009a); with RAD51 itself and with its paralogs RAD51C, RAD51AP1 and XRCC3 (Dray et al., 2010; Park et al., 2014a); with translesion polymerase η during recombination-associated DNA synthesis (Buisson et al., 2014); with KEAP1, an oxidative stress response protein (Ma et al., 2012); and with RNF168 ubiquitin ligase (Luijsterburg et al., 2017). PALB2 is ubiquitylated in G1 phase of the cell cycle by KEAP1 and CUL3, leading to its degradation, and, thereby, restraining its activity in S/G2 (Orthwein et al., 2015).

PALB2 promotes assembly of RAD51-ssDNA presynaptic nucleofilaments and formation of D-loop even in the absence of BRCA2 (Buisson et al., 2010; Dray et al., 2010). PALB2 recruits Polη polymerase to DSB sites and stimulates a recombination-associated DNA synthesis by Polη (Buisson et al., 2014).

BRCA1, −2 and PALB2 proteins also contain DNA binding domains (DBDs) (Buisson et al., 2010; Dray et al., 2010; Paull et al., 2001; Pellegrini et al., 2002). The functional role of DBDs in these proteins is poorly understood. Majority of missense mutations in the BRCA2 DBD are pathogenic (Guidugli et al., 2013; Wu et al., 2005). Disruption of BRCA2 DNA binding leads to HDR reduction with a BRCA2 construct lacking the PALB2-binding motif (Siaud et al., 2011). Recently, studies of BRCA1/BARD1 complex interaction with DNA and RAD51 led to the discovery of the BRCA1/BARD1 role in RAD51-mediated strand invasion and D-loop formation (Zhao et al., 2017). Two DBDs were previously identified in PALB2 (Buisson et al., 2010; Dray et al., 2010). The functional role of these domains remains unknown. PALB2 construct lacking 500 amino acids between the BRCA1 and BRCA2 binding motifs does not support BRCA2 and RAD51 foci formation in cells during DNA damage (Sy et al., 2009b). Since both the BRCA1-binding N-terminal and the BRCA2-interacting WD40 C-terminal domains were retained in this mutant, the results points to the potential importance of DBDs in PALB2 function.

In the current study, we identified a major DBD of PALB2 (N-DBD) and specific amino acids involved in DNA binding. Mutations of four amino acids significantly reduce RAD51 foci formation and the efficiency of HDR in a model cell system. Surprisingly, we found that N-DBD supports both forward and inverse strand exchange even in the absence of RAD51 and can use RNA as a substrate. Altogether, our data reveal a novel activity of PALB2 and highlight the importance of PALB2 DNA binding in chromosome maintenance in cells.

## RESULTS

### The DNA-binding mechanism of PALB2 and its function in DNA repair

#### The major DNA-binding site of PALB2 is localized in the N-terminal domain (N-DBD)

Two truncation fragments of PALB2 were previously reported to interact with DNA, T1 (residues 1-200) and T3 (residues 372-561)(Buisson et al., 2010). Both fragments together with the fragment consisting of amino acids 1-573 (PB2-573 in text), which includes both the T1 and T3, were cloned and purified (Fig. S1).

Quantitative measurement of PALB2 interaction with ss- and dsDNA oligonucleotides of different lengths demonstrate that T1 fragment alone interacts with all tested substrates with almost indistinguishable affinity from that of PB2-573 (Fig. 1). The T3 fragment has significantly lower affinity for DNA by itself. The K_d_ of the T1 and PB2-573 fragments were similar with both ss- and dsDNA substrates. The only difference was observed at an elevated salt concentration of 250 mM NaCl, where the PB2-573 fragment retained partial DNA binding activity (Fig. S2). In both cases, interactions were inhibited by addition of 500 mM NaCl. The T1 fragment will be referred as N-DBD in the text below. Interestingly, N-DBD binds long ssDNA substrates (49 nt) with significantly higher affinity than short ones (20 nt). This suggests an interaction with ssDNA through multiple binding sites, potentially formed by the previously described PALB2 oligomerization (Buisson and Masson, 2012; Sy et al., 2009c) or through interaction with multiple binding sites within a monomer (see below). Interaction with dsDNA was length-independent, suggesting that more rigid dsDNA interacts with a single site.

**Figure 1.**
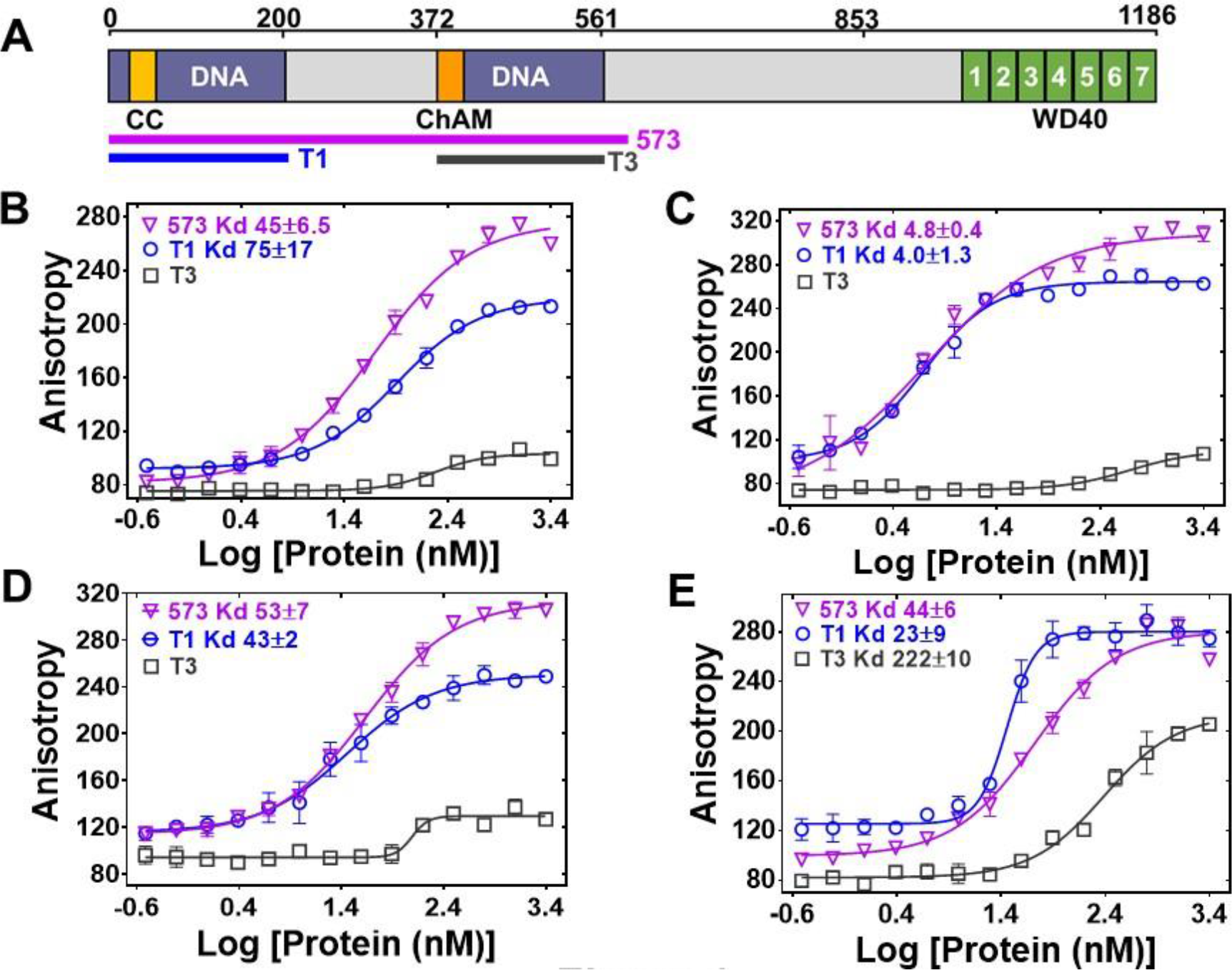
DNA binding affinity of T1, T3 and PB2-573 fragments to ss- and dsDNA. **(A)** Domain structure of PALB2. The PALB2 truncations used in the present study are shown below by magenta, blue and dark grey lines. **(B-E)** Equilibrium binding of PALB2 fragments, including T1 (blue), T3 (dark grey) and PB2-573 (magenta), to 20 nt ssDNA (ss20) **(B)**, 49 nt ssDNA (ss49) **(C)**, 20 bp dsDNA (ds20) **(D)**, and 49 bp dsDNA (ds49) **(E)** monitored by fluorescence anisotropy of FAM-labelled ssDNA. Each data point is an average of six readings from two different experiments. Inserts – binding parameters from nonlinear data fitting of titration curves for the T1 and 573 fragments. Reactions were performed in assay buffer with 20 mM Tris Acetate pH 7.0, 100 mM NaCl, 5% Glycerol, 10% DMSO 111 in a 40 μL reaction volume.

#### Identification of DNA-binding residues

Since the PALB2 DNA binding is salt dependent, we performed alanine scanning mutagenesis of several clusters of positively charged amino acids to identify the DNA binding site in the N-DBD (Fig. S3).

The main DNA-binding cluster is formed by amino-acids R146, R147, K148, and K149. Alanine mutation of these residues reduced binding affinity by two orders of magnitude with a K_d_ change from 4.0±1.3 nM to 316±59 nM in the case of T1 and from 4.8±0.4 nM to 167±50 nM in the case of PB2-573 (Fig. 2). DNA binding was moderately affected by mutations of two other clusters, including K45A/K50A, for which the K_d_ was increased to 28±5.2 nM (Fig. S4), and the triple mutant R170A/K174A/R175A with similar change in K_d_. From these experiments, we concluded that the main DNA binding site is formed by residues 146-149 with a potential minor contribution from other basic amino acids of the N-DBD.

**Figure 2.**
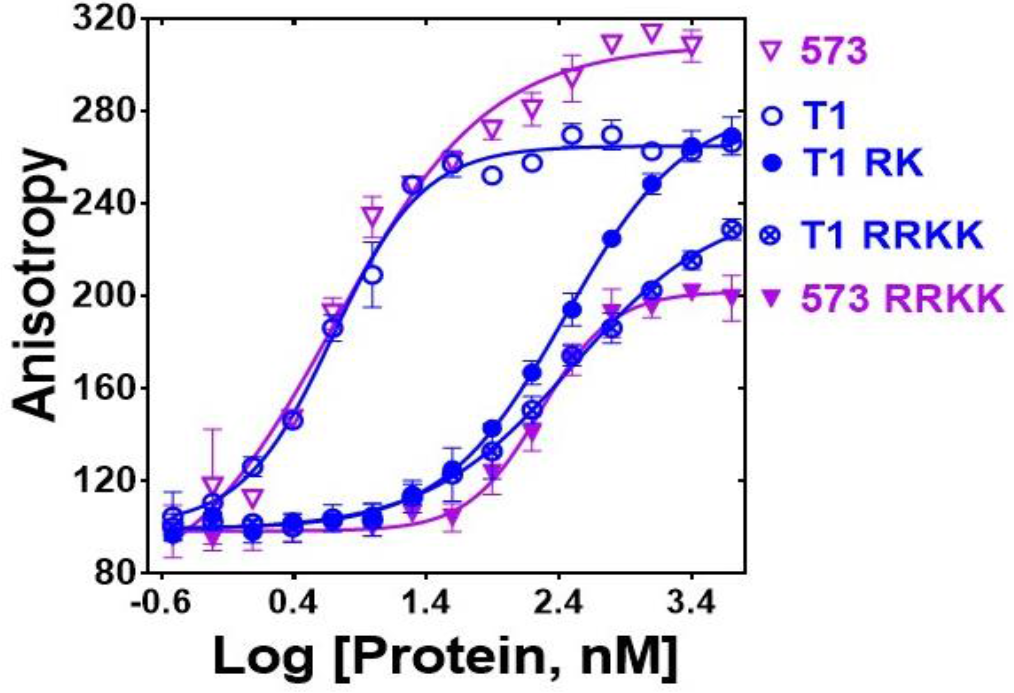
Mutation of DNA binding residues. Isotherm of fluorescence anisotropy of FAM-ss49 (5 nM) titrated by PALB2 T1 (blue, open circles) and PB2-573 (magenta, open triangles) fragments and their mutants: T1 146-RK/AA (filled blue circles), T1 146-RRKK/AAAA (crossed open blue circles), and 573 146-RRKK/AAAA (filled magenta triangles) under conditions identical to those in Fig. 1.

#### Impairment of DNA repair in cells with the PALB2 DNA-binding mutant

The mutations described above were used to separate the DNA-binding function from other macromolecular interactions of PALB2 during DNA repair in HeLa cell. Positively charged residues 146-149 were mutated to alanines in the full length PALB2 protein and the effect of these mutations was measured in two assays. First, we evaluated RAD51 foci formation in cells after gamma irradiation (Fig. 3A). Endogenous PALB2 was depleted by siRNA and cells were transformed with either wild type PALB2 or the DNA-binding mutant (Fig. 3A, bottom panel). PALB2 depletion leads to a severe defect in RAD51 foci formation. WT PALB2 restores RAD51 foci formation, while the DNA-binding PALB2 mutant restores only ~ 50% of RAD51 foci formation. Therefore, mutagenesis of only four positively charged residues in PALB2 has a major effect on efficiency of RAD51 recruitment to DNA damage sites.

Similarly, we tested the role of PALB2 interaction with DNA for the efficiency of HDR in U2OS cells using a novel LMNA-Clover based assay, where DNA breaks at a specific gene are introduced by the CRISPR system (Fig. S5) (Buisson et al., 2017b). As in case of RAD51 foci formation, complementation of PALB2-depleted cells with the DNA-binding PALB2 mutant restores only 50% of HDR efficiency, in contrast to WT PALB2, which restores more than 90% of activity (Fig. 3B). Altogether, these studies show that PALB2 DNA binding plays a significant role in HR and DNA repair *in vivo*.

**Figure 3.**
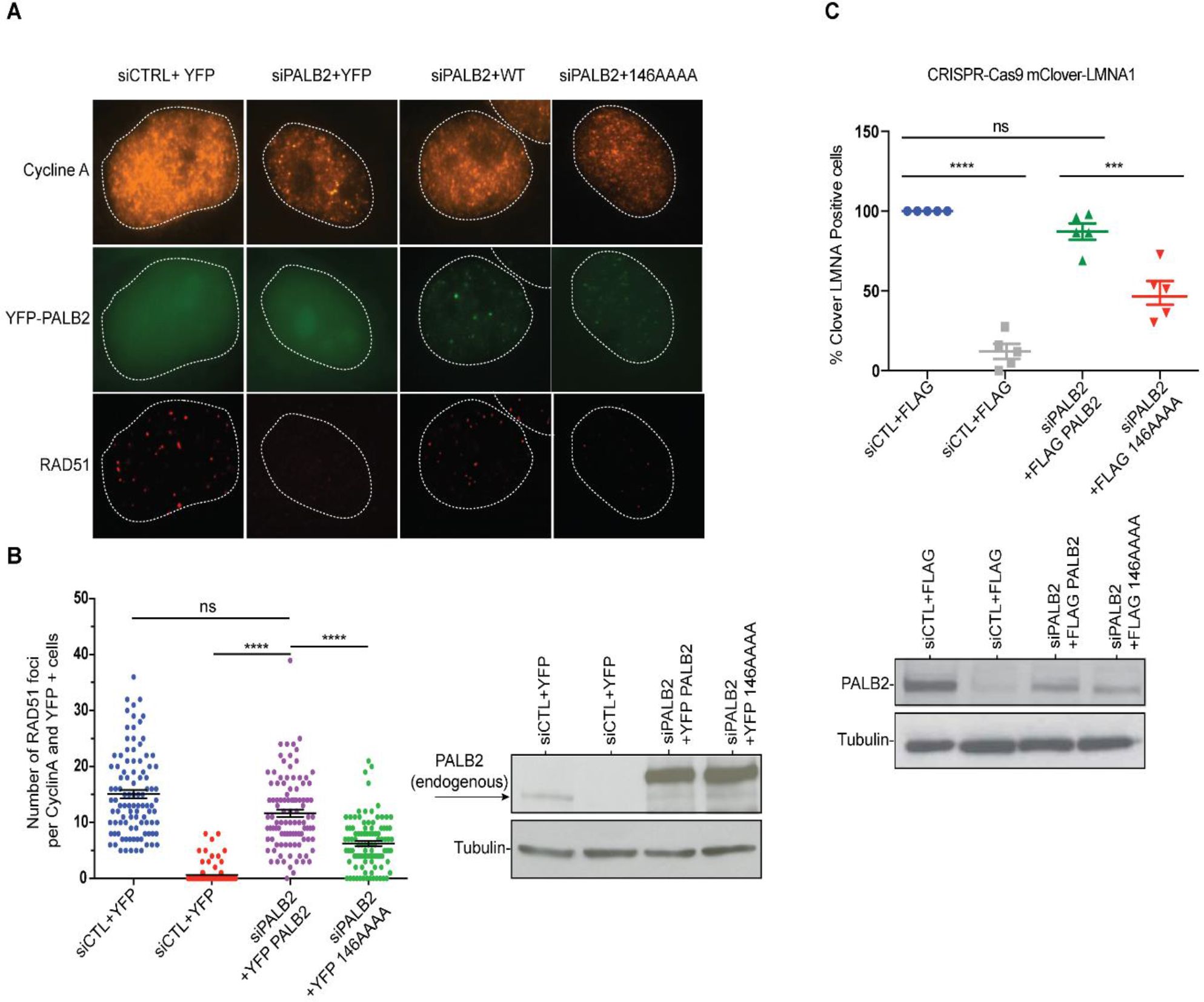
Effect of a PALB2 DNA-binding mutation on homologous recombination. **A)** Representative immunofluorescence images of RAD51 foci in PALB2 knockdown HeLa cells complemented with the indicated YFP construct and synchronized in S/G2 by double thymidine block, as determined by cyclin A staining. **B)** Left: RAD51 foci quantification in control siRNA (blue), siPALB2 (red) and with siPALB2 with subsequent complementation by siRNA resistant constructs YFP-PALB2 (magenta) and 146AAAA DNA-binding mutant PALB2 (green). Right: Western blotting of the samples shown in B to monitor knockdown and complementation efficiency. **C)** Top: Gene-targeting efficiency of siRNA PALB2 cells complemented with wild-type and 146AAA siRNA resistant constructs mClover positive/iRFP cells were quantified. Bottom: Western blotting of the samples shown in C) to monitor knockdown and complementation efficiency. ***P<0.01 and ****P<0.001. (Fig S5B).

### PALB2 promotes DNA and RNA strand exchange

#### PALB2 stimulates RAD51-mediated strand exchange and promotes a similar reaction without RAD51

PALB2 stimulates RAD51 filament formation even in the absence of BRCA2 (Buisson et al., 2010). Here, we investigated the ability of the PALB2 N-DBD to stimulate the strand exchange activity of RAD51 using a fluorescence-based strand exchange assay similar to the one previously published (Fig. 4A) (Jensen et al., 2010; Ryzhikov et al., 2014). Under solution conditions used in DNA-binding assays in Fig. 1 and even with reduced NaCl concentration, RAD51 displayed a low activity, in contrast to *E. coli* RecA (Fig. 4B, C).

**Figure 4.**
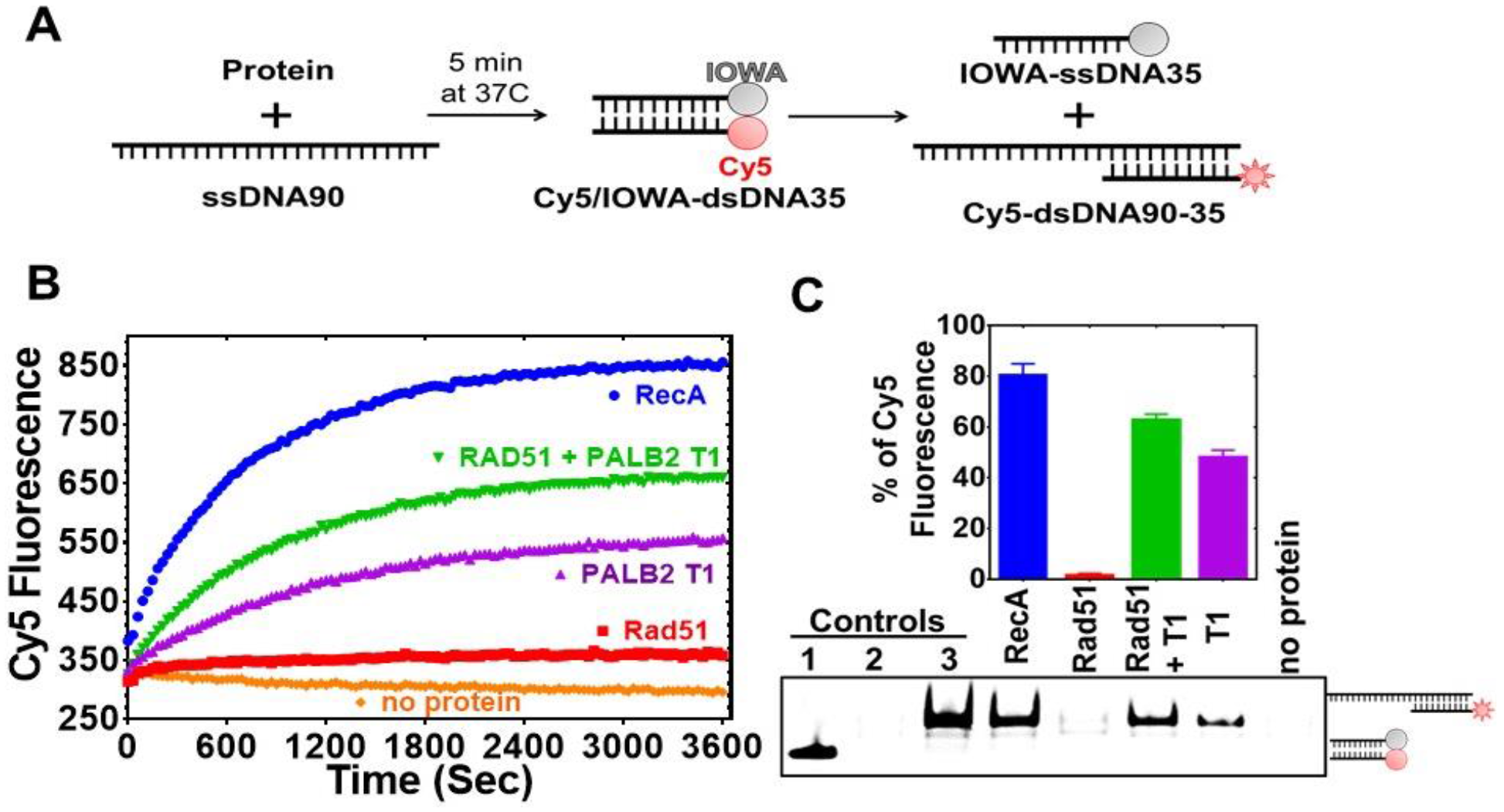
PALB2 promotes strand exchange between homologous DNA substrates. **A)** Schematic representation of the strand exchange activity assay. ss90 was incubated with RecA (2μM) or RAD51 (2 μM) for 5’, then with PALB2 fragment (2 μM) for 5’, then dsDNA was added and the Cy5 fluorescence was measured on a plate reader using 635 nm excitation and 680 nm emission wave-lengths. **B)** Continuously measured Cy5 fluorescence after initiating reactions with RecA (blue), RAD51 (red), PALB2 N-DBD (magenta), RAD51 and PALB2 N-DBD (green), and without proteins (orange). **C)** Reaction products from B) were deproteinized and separated on a native PAGE gel. Bar graph represents the percentage intensities of strand exchange products on the gel relative to the intensity of control product (Cy5-dsDNA35-90).

RAD51 activity was stimulated by addition of 5 mM CaCl_2_ (Fig. S6). Recombination mediator proteins (RMPs) stimulate recombinase activity even at unfavourable solution conditions, such as in the case of Rad52 (Krejci et al., 2002; New et al., 1998), BRCA2 (Jensen et al., 2010)(Liu et al., 2010; Thorslund et al., 2010) and the Hop2-Mnd1 complex (Chi et al., 2007). Similar to the previously published finding that the full length PALB2 stimulates RAD51 function (Buisson et al., 2010; Dray et al., 2010), we found that the PALB2 N-DBD alone stimulates RAD51-mediated strand exchange (Fig. 4).

Surprisingly, the N-DBD promotes strand exchange at a comparable rate even without RAD51. Reaction products were further analysed by EMSA gel shift to rule out any artefact of protein-specific fluorophore quenching (Fig. 4C). The results were confirmed using DNA with different fluorescent labels (Fig. S7). The strand exchange activity of N-DBD was even more efficient with longer dsDNA substrate (Fig. S8).

Since N-DBD stimulates a similar reaction on its own, it is unclear whether the N-DBD fragment stimulates RAD51 activity or if the two proteins function independently. In a limited titration experiment shown in Figure 5, the efficiency of the strand exchange increases proportionally to N-DBD concentration in the absence of RAD51. However, in the presence of 2 μM of RAD51, the maximum rate of strand exchange is reached at 1 μM of N-DBD. These data suggest a synergistic effect of two proteins in a strand exchange reaction.

**Figure 5.**
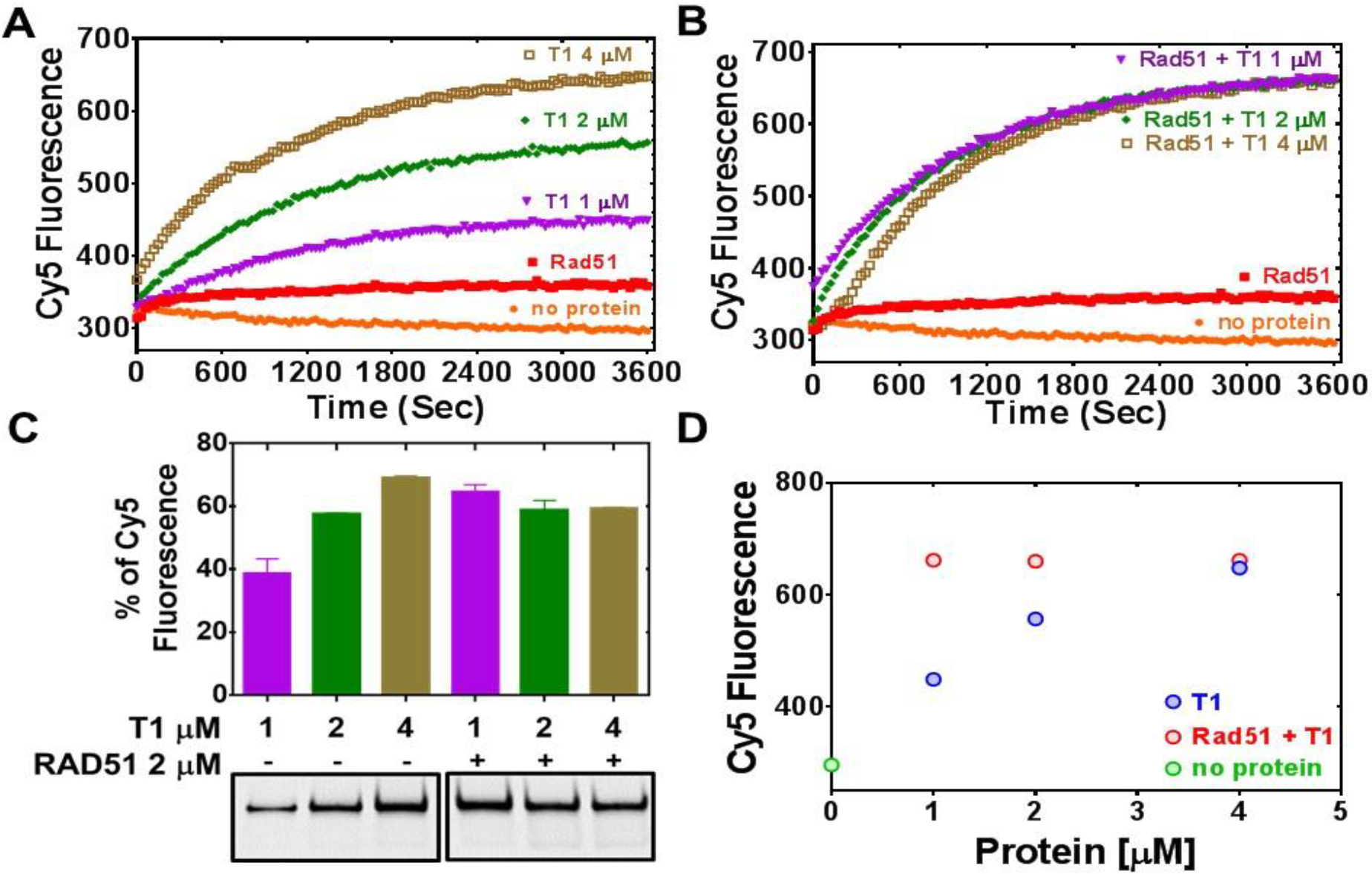
The PALB2 N-DBD stimulates RAD51 strand exchange. **A)** Strand exchange activity of PALB2 with increasing concentration of N-DBD: 1 μM (magenta), 2 μM (green), and 4 μM (light brown). **B)** Stimulation of RAD51 (2 μM, red) strand exchange activity by PALB2 N-DBD at the same concentrations. **C)** Deproteinated strand exchange activity products from A) and B) separated on native PAGE gel. **D)** End-point values of the efficiency of strand exchange reactions shown in A) and B) plotted against N-DBD concentration.

Additional evidence of synergism between the two proteins comes from conformational changes of ssDNA labelled with a Cy3 at 5’ and Cy5 at 3’ ends (Cy3-dT_70_-Cy5) (Figs. 6, S9). PALB2 N-DBD supports a compact conformation with a high FRET, as in the cases of RAD52 (Grimme et al., 2010) and SSB (Roy et al., 2007). RAD51 binds ssDNA in an extended confirmation with a low FRET. Both proteins support an intermediate FRET value that remains constant even with changed molar ratio of two proteins in solution. The conformation became more extended upon addition of ATP in the case of RAD51 and of RAD51 with N-DBD (Fig. S9). Addition of excess N-DBD does not increase FRET, suggesting that RAD51 and N-DBD form a stronger complex with ssDNA than N-DBD alone. Both experiments together (Figs. 5, 6, S9) strongly support formation of a complex between PALB2 N-DBD and RAD51 on ssDNA and synergism during the presynaptic filament formation and the strand exchange.

**Figure 6.**
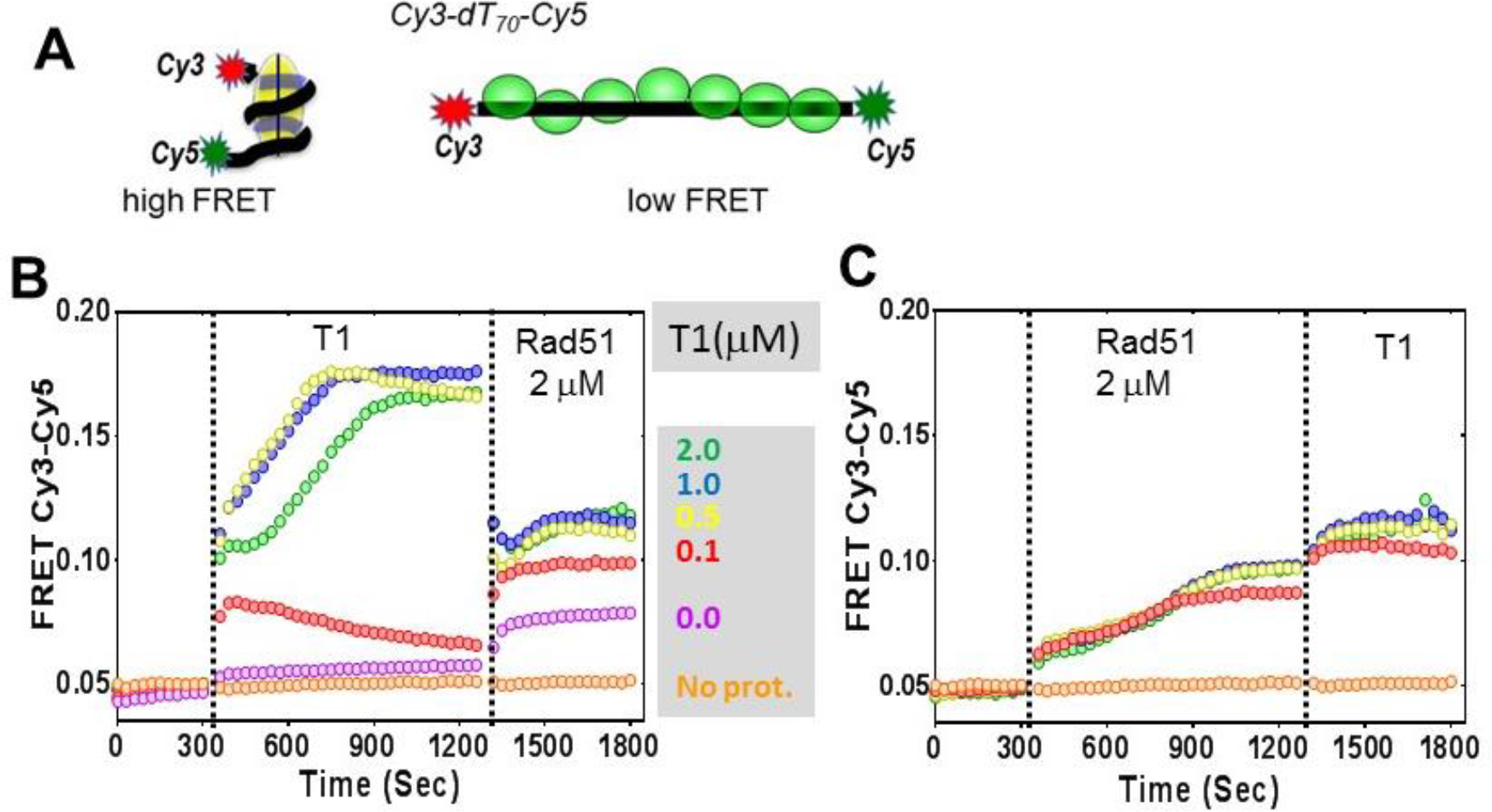
Cooperative interaction of PALB2 and RAD51 with ssDNA. **A)** Schematic representation of the experiment showing Cy3 and Cy5 labelled ssDNA in compact conformation wrapped around the protein (left) or in extended conformation supported by the recombinase filament (right). **B)** FRET values of Cy3-dT_70_-Cy5 alone (initial 5 min) and upon addition of different amounts of N-DBD (color-coded) and then by addition of 2 μM of RAD51 in the absence of ATP. **C)** Similar experiment but with RAD51 added first followed by the addition of the PALB2 N-DBD. Similar experiment conducted in the presence of ATP is shown in Fig. S9.

#### PALB2 stimulates an inverse strand exchange and can use an RNA substrate

RecA and Rad52 support an inverse strand exchange as well as an R-loop formation (Kasahara et al., 2000; Mazina et al., 2017; Zaitsev and Kowalczykowski, 2000). We tested the PALB2 N-DBD for similar activities. The PALB2 N-DBD supported both forward and inverse strand exchange with similar efficiencies (Fig. 7B, E). Furthermore, PALB2 supported both reactions with a ssRNA substrate (Figs. 7C, F). DNA-binding mutant fragment (146AAAA) did not support strand exchange on its own and in the presence of RAD51 (Fig. S10). RAD52 was shown to have different efficiency of forward and inverse reactions with relatively low forward and a more efficient inverse reactions (Mazina et al., 2017). We did not observe this difference with PALB2. The inverse strand exchange was slower than in case of RAD52 and comparable to that of RAD51 under optimal conditions.

**Figure 7.**
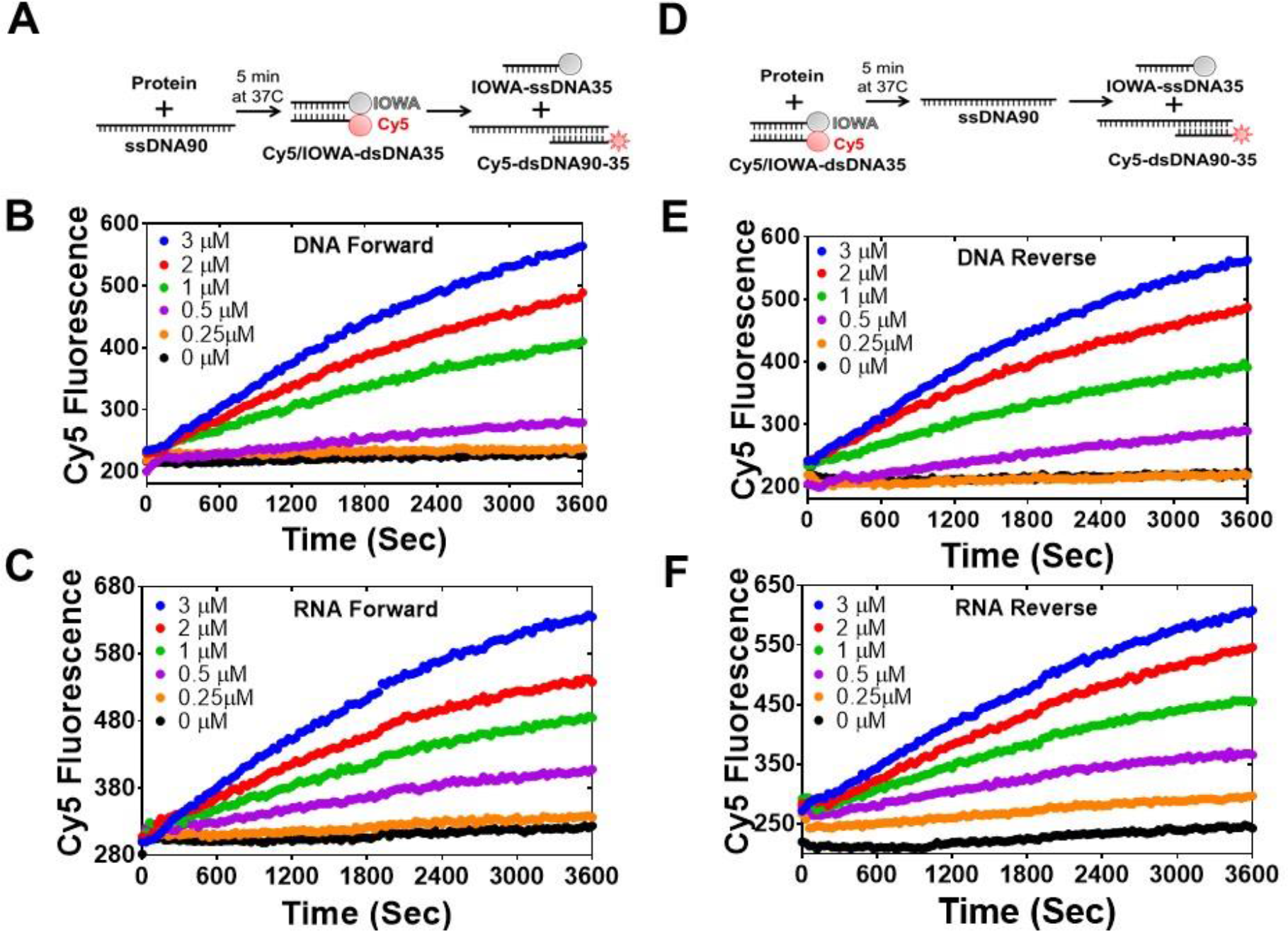
PALB2 promotes forward and inverse strand exchange with ssDNA and ssRNA substrates. Schematic representation of forward **(A)** and inverse **(D)** reactions. Cy5 fluorescence change for the forward reaction with ssDNA is shown in **(B)** and with ssRNA in **(C)** at different concentrations of T1 fragment ranging from 3 μM (blue) to 0.25 μM (orange) and without protein in black. Time courses of inverse reaction are shown in (**E)** for ssDNA and in **(F)** for ssRNA substrates, correspondingly.

#### Mechanism of the PALB2 stimulated strand exchange

To rule out a potential effect of DNA melting by PALB2, which may lead to nonspecific reannealing of a separated strands with complementary ssDNA in solution, the N-DBD was incubated with dsDNA without ssDNA (Fig. S11). The N-DBD does not melt dsDNA as there was no change in fluorescence of Cy5/Iowa-ds35 upon incubation with the protein, while the addition of complimentary ssDNA triggers the reaction. Moreover, the N-DBD stimulates annealing of complimentary ssDNA (Fig. S12). Therefore, the observed strand exchange is is not a consequence of a nonspecific dsDNA melting by the protein.

Both RecA and RAD52 proteins, which support strand exchange, simultaneously interact with ds- and ssDNA through distinct binding sites located next to each other (Arai et al., 2011; Chen et al., 2008; Honda et al., 2011; Kagawa et al., 2002; Mazin and Kowalczykowski, 1998; Seong et al., 2008). PALB2 also interacts with both ss- and dsDNA (Fig. 1 and Buisson et al., 2010; Dray et al., 2010), although the structure and the molecular details of PALB2 interaction with DNA remains unknown. To verify if other proteins characterized by comparable affinities to both ss- and dsDNA also support strand exchange, we tested prokaryotic RMPs, RecO and RecR. *E. coli* RecO alone stimulates strand annealing (Kantake et al., 2002; Luisi-DeLuca and Kolodner, 1994) and, in complex with RecR, stimulates RecA-mediated strand exchange with ssDNA bound to SSB (Ryzhikov et al., 2014; Umezu et al., 1993; Umezu and Kolodner, 1994). Both RecO and RecOR interact with ss- and dsDNA (Ryzhikov et al., 2014). However, neither RecO nor RecOR complex promote strand exchange in the absence of RecA (Fig. S13). Therefore, the simple ability of a protein to interact with ss- and dsDNA is not enough to promote strand exchange and even RMPs, which stimulate the reaction by RecA recombinase, do not support it in the absence of recombinase.

## DISCUSSION

In this report, we identify major DNA-binding residues of PALB2 and demonstrate their critical role for the HDR in cells. PALB2 is described as a scaffold protein linking BRCA1 with BRCA2 during HDR and interacting with many other chromatin proteins. However, the mutant with BRCA1 and BRCA2 binding motifs without the middle portion of the proteins does not support BRCA2 and RAD51 recruitment to DSBs (Sy et al., 2009b). A critical role of PALB2 DNA binding was also suggested by studies of the BRCA2 mechanism (Siaud et al., 2011), where the “miniBRCA2” construct, which includes only DBDs with two BRC repeats, was 3-4 times less efficient in the absence of PALB2 interaction. Moreover, interaction with PALB2 alleviates the requirement of BRCA2 DNA binding, including a deletion of the entire BRCA2 DBD. Here, we demonstrate that mutation of only four DNA-binding residues of PALB2 reduces both RAD51 foci and overall HDR efficiency by 50%, even in the presence of endogenous BRCA2. Therefore, PALB2 interaction with DNA is critical for recruitment of the BRCA2 and RAD51 to DSB sites and efficient DNA repair in cells.

Secondly, we demonstrate that PALB2 N-DBD stimulates RAD51-mediated strand exchange *in vitro*. Therefore, PALB2 can cooperate with BRCA2 in loading RAD51 onto DNA and/or promoting the subsequent steps of D-loop formation and recombination dependent DNA synthesis.

The most unexpected finding is the ability of PALB2 to stimulate strand exchange between homologous ss- and dsDNA fragments in the absence of recombinase. This process is protein specific and is not a consequence of a simple DNA melting and reannealing of separated strands in solution, since the PALB2 N-DBD does not unwind the DNA helix and promotes DNA annealing. Proteins supporting strand exchange, such as RecA and RAD52, share several common features. They interact with both ss- and dsDNA through distinct sites located next to each other (Arai et al., 2011; Chen et al., 2008; Honda et al., 2011; Kagawa et al., 2002; Mazin and Kowalczykowski, 1998; Saotome et al., 2018; Seong et al., 2008), they form oligomeric structures, such as recombinase-DNA filament (Chen et al., 2008; Egelman and Stasiak, 1986; Yang et al., 2001) or Rad52 ring structure (Shinohara et al., 1998; Singleton et al., 2002), and they distort the dsDNA helix to initiate strand exchange with the bound complementary ssDNA. RecA stretches dsDNA (Chen et al., 2008; Leger et al., 1998), while Rad52 bends the DNA helix bound to the toroidal oligomeric ring (Brouwer et al., 2017). The PALB2 N-DBD interacts with both ss- and dsDNA. Both N-DBD and the full length PALB2 form oligomeric structures and the oligomerization is partially mediated by the N-terminal coiled-coil motif (Fig. S14A) and (Buisson and Masson, 2012; Sy et al., 2009c). Titration of ss49 by the N-DBD suggests a stoichiometry of four or five N-DBD monomers per ss49 (Figs. S14B, C) which fits an oligomeric state suggested by the size-exclusion chromatography data (Fig. S14A).

Previously, we demonstrated that PALB2 immobilized on ssDNA beads efficiently pulls down non-homologous dsDNA (Buisson et al., 2010). It is unclear whether ss- and dsDNA substrates are bound to same site of different subunits of PALB2 oligomer of two different DNA binding site of the same monomer. The presence of at least two other minor DNA-binding sites in the N-DBD suggests such a possibility. Higher affinity towards longer ssDNA and the FRET experiment support a model of wrapping long flexible ssDNA around an oligomer and binding to multiple sites. In contrast, interaction with dsDNA is less length dependent. We can speculate that binding of dsDNA to more than one monomer in PALB2 oligomer can trigger DNA helix distortion. Indeed, bending of dsDNA was observed upon PALB2 binding to 40 bp dsDNA labelled with a Cy3/Cy5 FRET pair as the increase of FRET signal (Fig. S15). Thus, PALB2 shares several specific structural and DNA interaction features with both RecA/RAD51 and Rad52 proteins and supports a protein-specific strand exchange reaction.

It is important to note one distinct feature of the PALB2 N-DBD: the secondary structure prediction (Fig.S3) suggests a different folding of the N-DBD fragment than that of RecA-like domains or a Rad52. The latter proteins are formed by α/β sandwich folds, while PALB2 N-DBD folding is predicted to be composed of only α-helices, which may form helical bundle-like structure similar to that of Hop2-Mnd1(Kang et al., 2015). Therefore, PALB2 N-DBD represents a novel structural fold that supports strand exchange.

While the functional significance of this property remains to be investigated, it further supports the involvement of PALB2 in specific DNA transactions during HDR, similar to the involvement of BRCA1/BARD1 in D-loop formation (Zhao et al., 2017). It was shown that both PALB2 and BRCA2 stimulate Polη DNA synthesis within a D-loop substrate *in vitro* through the recruitment of the polymerase to the invading strand in the D-loop (Buisson et al., 2014). Interestingly, DNA synthesis was more efficient in the presence of PALB2 than BRCA2, while both proteins were shown be equally efficient in recruiting polymerase to DSB sites. PALB2 strand exchange may contribute to other steps of HDR such as second-end capture(Mazloum and Holloman, 2009; McIlwraith and West, 2008; Nimonkar et al., 2009). Interestingly, PALB2 (FANCN) and BRCA2 (FANCD) are involved in replication-dependant removal of interstrand DNA crosslinks associated with Fanconi anemia (Howlett et al., 2002; Moldovan and D’Andrea, 2009; Xia et al., 2007). The strand exchange function of PALB2 may be also important for alternative DNA repair pathways. Indeed, PALB2 supports strand exchange not only with ssDNA, but with ssRNA substrates, and can be involved in transcription-initiated DNA repair.

## MATERIAL AND METHODS

### Protein Purification

#### PALB2 truncations

PALB2 N-terminal fragments PALB2-T1 (1-200 aa) and PALB2-573 (1-573 aa) were cloned into pET28b+ based pSMT3 vector (provided by Dr. R.A. Kovall, University of Cincinnati) containing the N-terminal 6xHis-SUMO tag using *Sal*I and *Nde*I cloning sites. pSMT3-PALB2 T1, pSMT3-PALB2 573 were transformed into BL21* cells. Cell culture were grown in LB to OD600=0.7 and protein expression was induced by adding 0.2 mM IPTG and carried out at 16°C overnight. Cells were lysed with lysozyme (0.25 mg/mL at RT for 30 min) in lysis buffer (25 mM HEPES pH 8.0, 1 M NaCl, 10% Glycerol, 0.3% Brij35, 1 mM TCEP, 2 mM CHAPS and 1 mM PMSF), followed by three rounds of sonication (50% output and 50% pulsar settings for 4 min). Cell debris was removed by centrifugation at 30,600 x g for 45 min. Supernatant was loaded on a NiNTA column (5 ml) equilibrated with binding buffer (25 mM HEPES pH 8.0, 1 M NaCl, 10% glycerol, 1 mM TCEP, 2 mM CHAPS and 10 mM imidazole). NiNTA beads were washed with binding buffer and the protein was eluted with binding buffer supplemented with the same buffer adjusted to 250 mM imidazole. The SUMO tag was cleaved with Ulp1 protease while dialyzing against buffer without imidazole (25 mM HEPES pH 8.0, 1 M NaCl, 10% Glycerol, 1 mM TCEP and 2 mM CHAPS) overnight and the protein was purified with a second NiNTA column. The protein was diluted 10X by binding buffer without NaCl to the final NaCl concertation of 100 mM, loaded to a Hi-Trap heparin affinity column (5 ml, GE health sciences) and eluted with a gradient of NaCl (100 mM to 1000 mM). Protein eluted from the heparin column at ~500 mM NaCl concentration. Protein fractions were dialysed against storage buffer (25 mM HEPES pH 8.0, 300 M NaCl, 40% Glycerol, 1 mM TCEP and 2 mM CHAPS) overnight, aliquoted and stored in − 80°C.

PALB2 T3 fragment was purified as described in (Buisson et al., 2010).

#### RAD51 purification

We used two expression constructs and purification protocols. 1) Human RAD51 protein was purified from the pET11-Rad51 vector (gift from Dr A. Mazin) similarly to the published protocol (Sigurdsson et al., 2001). The protein was induced at 37°C for 3h by supplementing LB media with 0.5 mM IPTG. Cells were suspended in 25 mM Tris-HCl pH8.0, 1 M urea, 1 M NaCl, 5 mM DTT, 0.3 % Brij35 and 10 % glycerol. Cells were lysed with lysozyme (0.25 mg/mL at RT for 30 min) followed by three rounds of sonication (50% output and 50% pulsar settings for 1 min). 24 mg/ml ammonium sulphate was gradually added to the supernatant and equilibrated overnight at 4°C. Precipitates were centrifugation at 30,600 x g for 45 min. Pellets were solubilized in 30 ml of binding buffer (25 mM Tris-HCl pH8.0, 1 M NaCl, 5 mM DTT, 10 % glycerol and 20 mM imidazole). Insoluble particles were removed by centrifugation at 30,600 x g for 40 min. The protein was bound to Ni NTA beads, extensively washed with binding buffer and eluted in binding buffer supplemented with 250 mM imidazole.

2) Alternatively, human RAD51 gene was cloned into pSMT3 vector using *Sal*I and *Nde*I cloning sites. pSMT3-Rad51 protein expression was carried out at 16°C overnight by addition of 0.2 mM IPTG. SUMO tagged Rad51 protein was purified according to the steps described for the PALB2 fragments. Purified Rad51 protein was dialysed against storage buffer (25 mM HEPES pH 8.0, 300 M NaCl, 40% glycerol, 1 mM TCEP and 2 mM CHAPS) overnight, aliquoted and stored in −80°C until further use. Proteins from both preparations had comparable properties. Data are shown for experiments performed with the second construct, except the experiment represented in Fig. S10.

*E. coli* RecA was purified exactly as described in (Gupta et al., 2013). *E.coli* RecO and RecR proteins were purified as described in (Ryzhikov and Korolev, 2012; Ryzhikov et al., 2011).

### Site-directed mutagenesis

Target amino acids were mutated by site directed mutagenesis of using Stratagene QuikChange™ protocol. Single, double, triple and four residues mutants were generated by single stranded synthesFis (Table S1). PCR samples were subjected to *Dpn*I digestion at 37°C for 6 h and annealed gradually by reducing temperature from 95°C to 37°C for an hour with a degree 1C drop per min. *Dpn*I treated PCR samples were transformed into chemically competent OmniMAX cells (ThermoFischer). Mutations were confirmed by sequencing and plasmids were transformed into BL21(DE) cells. The PALB2 573 AAAA mutant was generated as described for PALB2 T1. Mutant proteins were expressed and purified exactly as described for wild type fragments.

### DNA binding assay

Fluorescence anisotropy experiments were carried out at room temperature with 5 nM fluorescein (6FAM)-labelled DNA substrates (Table S2) using a Synergy 4 plate reader (BioTek). Titration with protein was performed by serially diluting protein in 40 μL of assay buffer (20 mM Tris Acetate pH 7.0, 100 mM NaCl, 5 % glycerol, 1 mM TCEP and 10% DMSO) from 5000 nM to 0.3 nM and incubating with DNA substrate for 15 min at RT. Fluorescence anisotropy was measured by excitation at 485/20 nm and by monitoring emission 528/20 nm at room temperature. Data were fit using a standard four-parameter logistic fit (Prism).

### DNA annealing assay

DNA annealing assays were performed with Cy5-labelled ss35 and complimentary IOWA-labelled ss35 (100 nM, Table S3). The protein at 1, 2 and 4 μM concentrations was mixed and incubated with Cy5-labeled ssDNA for 5 min at 37°C in 40 μL reaction buffer (40 mM HEPES pH 7.5, 20 mM NaCl and 1 mM TCEP). Reaction was initiated by addition of complimentary IOWA-labelled ss35 (100 nM) in 40 μL of reaction buffer. Decrease in Cy5 fluorescence was monitored by measuring fluorescence at 680 nm by excitation at 635nm on a Synergy 4 plate reader (BioTek).

### Strand exchange fluorescent assays

DNA strand exchange assays (80 μl) were performed with 35bp dsDNA obtained by annealing of 5’-Cy5- and 3’-IOWA labelled complimentary strands (Table S3) and a 90mer ssDNA (ss90) with homologous region to plus strand. Alternatively, FAM/Dabsyl 49 bp DNA was used. For the forward reaction, ss90 (100 nM) was incubated with 2 μM (or as mentioned in the figure legends) protein for 10 min in 40 μL reaction buffer (40 mM HEPES pH 7.5, 20 mM NaCl, 5 mM MgCl_2_, 1 mM TCEP and 0.02 % Tween 20) at 37°C. Strand exchange was initiated by addition of 100 nM 35bp dsDNA (ds35), plate was immediately placed in plate reader and the intensity of Cy5 fluorescence was measured at 30 sec intervals for 1 hour with excitation at 635 nm and emission at 680 nm. For reactions with RecA and Rad51, an ATP regeneration system (2 mM ATP, 30 mM phosphoenol pyruvate and 30 U of pyruvate kinase) was used (Sigma-Aldrich, USA). For the inverse reaction, protein was incubated with Cy5/IOWA-dsDNA35 substrate and reaction was initiated by addition of ss90. The strand exchange assay with an ssRNA substrate was performed as described above using a 60 ribonucleotide RNA (table S1) complimentary to that of 35bp DNA. Alternatively, Cy3- and Cy5-labelled DNA oligonucleotides were used to prepare dsDNA substrate and the products were analysed by EMSA PAGE (below).

### EMSA PAGE

Fluorescent-labelled DNA products of strand exchange reactions were also analysed on EMSA PAGE. After fluorescence measurement on plate reader, the final reaction mix (80 μl) products were deproteinated by incubation with proteinase K (0.5 mg/ml) with 0.5 mM EDTA and 1% w/v SDS for 20 min at 37°C and the DNA fragments were separated on 10% PAGE gel in TBE buffer. The gel was imaged using a Typhoon 9400 image scanner (GE) and analysed with ImageJ software.

### FRET assay

FRET assay was performed in 96 well plate format. 100 nM of dual labelled dT_70_ (Cy5 at 5’ end and Cy3 at 3’ end) was dispensed into 80 μL assay buffer identical to those in strand exchange assay (Table 3). Alternatively, dual labelled 40 bp DNA was prepared by annealing dual labelled 40 nt ssDNA (Cy3 and Cy5 on single strand) with unlabelled complimentary 40 nt ssDNA (Table 3). Excitation was at 540/25nm bandpass. Emission for both Cy3 at 590/35nm bandpass and Cy5 at 680/30 nm were monitored at 30 s intervals for 5 min at 37°C, then for 10 min following the addition of PALB2 N-DBD, then for 10 min following the addition of RAD51 with or without ATP. FRET efficiency was calculated by using the formula 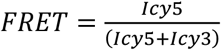. For each addition, the plate was removed from the plate reader and returned to the reader within 60 s. Protein concentrations were as described in the figure legends.

### RAD51 foci assay

HeLa cells were seeded on glass coverslips in 6-well plates at 225 000 cells per well. Knockdown of PALB2 was performed 18 hours later with 50 nM PALB2 siRNA (Table S4) using Lipofectamine RNAiMAX (Invitrogen). After 5 hours, cells were subjected to double thymidine block. Briefly, cells were treated with 2 mM thymidine for 18 hours and released after changing the media.

After a release of 9h, PALB2 silenced cells were complemented using transfection with the indicated YFP constructs using Lipofectamine 2000. Then, cells were treated with 2 mM thymidine for 17 hours and protected from light from this point on. After 2 h of release from the second block, cells were X-irradiated with 2 Gy and processed for immunofluorescence 4 h post-irradiation. All immunofluorescence dilutions were prepared in PBS and incubations performed at room temperature with intervening washes in PBS. Cell fixation was carried out by incubation with 4% paraformaldehyde for 10 min followed by 100% ice-cold methanol for 5 min at −20 °C. Cells were then permeabilized in 0.2% Triton X-100 for 5 min and quenched using 0.1% sodium borohydride for 5 min. After blocking for 1 h in a solution containing 10% goat serum and 1% BSA, cells were incubated for 1 h with primary antibodies to RAD51 (B-bridge International, #70-001) and to cyclin A (BD Biosciences, # 611268) diluted in 1% BSA. Secondary antibodies, Alexa Fluor 568 goat anti-rabbit (Invitrogen, #A-11011) and Alexa Fluor 647 goat anti-mouse (Invitrogen, #A-21235), were used in PBS containing 1% BSA for 1 h. Nuclei were stained for 10 min with 1 μg/mL 4, 6-diamidino-2-phenylindole (DAPI) prior to mounting onto slides with 90% glycerol containing 1 mg/ml paraphenylenediamine anti-fade reagent. Z-stack images were acquired at 63X magnification on a Leica DM6000 microscope, then deconvolved and analysed for RAD51 foci formation with Volocity software v6.0.1 (Perkin-Elmer Improvision). The number of RAD51 foci per cyclin A-positive cells (n=100), among the transfected population, was manually scored and reported in a scatter dot plot representing the SEM. An Anova test (Kruskul-Wallis test for multiple comparison) was performed followed by a non-parametric Mann-Whitney test.

### CRISPR Cas9/mClover-LMNA1 mediated HR assay (Pinder et al., 2015)

U2OS cells were seeded in 6-well plates. Knockdown of PALB2 (Buisson et al., 2017a) was performed 6-8 h later using Lipofectamine RNAiMAX (Invitrogen). Twenty-four hours post-transfection, 1.5-2×10^6^ cells were pelleted for each condition and resuspended in 100 μL complete nucleofector solution (SE Cell Line 4D-Nucleofector™ X Kit, Lonza) to which 1μg of pCR2.1-mClover-LMNAdonor, 1μg pX330-LMNAgRNA, 0.1μg of iRFP670 and 1μg of pcDNA3 empty vector or the Flag-PALB2 constructs, and 20nM of each siRNA were added. Once transferred to a 100 ul Lonza certified cuvette, cells were transfected using the 4D-Nucleofector X-unit, program CM-104 and transferred to a 10 cm dish. After 48 h, cells were trypsinized and plated onto glass coverslips. Expression of the mClover was assayed the next day by fluorescence microscopy (63X), that is 72h post-nucleofection. Data are represented as mean percentages of mClover-positive cells over the iRFP-positive population from five independent experiments (total n>100 iRFP-positive cells) and reported in a scatter dot plot representing SEM, and a classical one-way Anova test was performed.

### Plasmids and siRNA

peYFP-C1-PALB2 was modified to be resistant to PALB2 siRNA by Q5 Site-Directed Mutagenesis Kit (NEB, E0554) using primers JYM3892/3893 (Table S4). The resulting siRNA-resistant construct was then used as a template to generate the mutant construct YFP-PALB2 146AAAA with the primers JYM3909/JYM3910. Flag-tagged PALB2 146AAAA mutant was also obtained via site-directed mutagenesis on pcDNA3-Flag PALB2 (Pauty et al., 2017).

## SUPPLEMENTARY DATA

Supplementary data are available in Supplementary Figures.

## ACKNOWLEDGEMENTS

We are grateful to the members of Korolev lab including Ian Miller for help with cloning, Lakshmi Kanikkannan and Jennifer Redington for help with protein purification and DNA-binding assays. We are grateful to Drs Alessandro Vindigni and Joel Eissenberg for the manuscript evaluation and discussions.

---

**Table S1.**
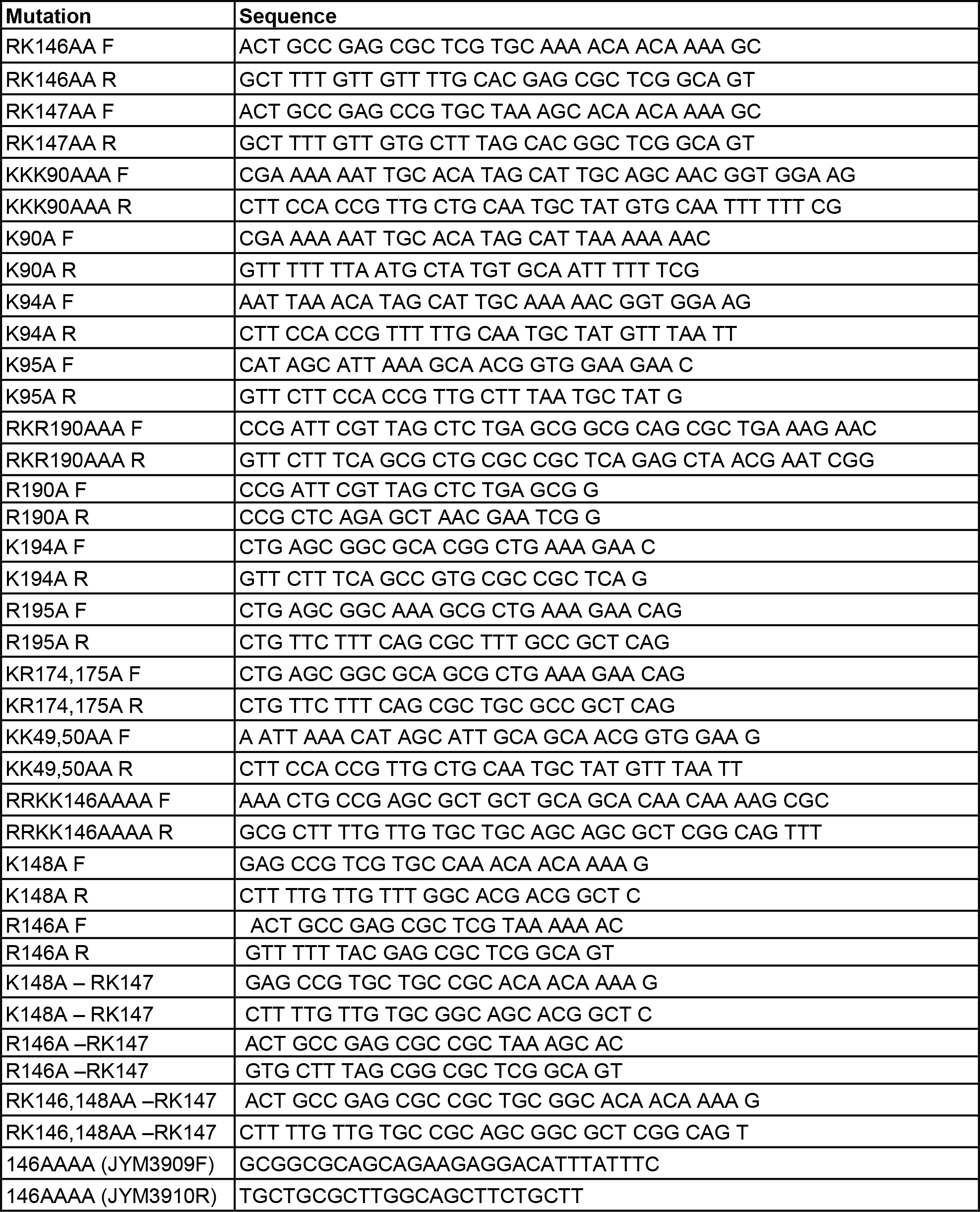
Primers for DNA binding site mutagenesis.

**Table S2.**
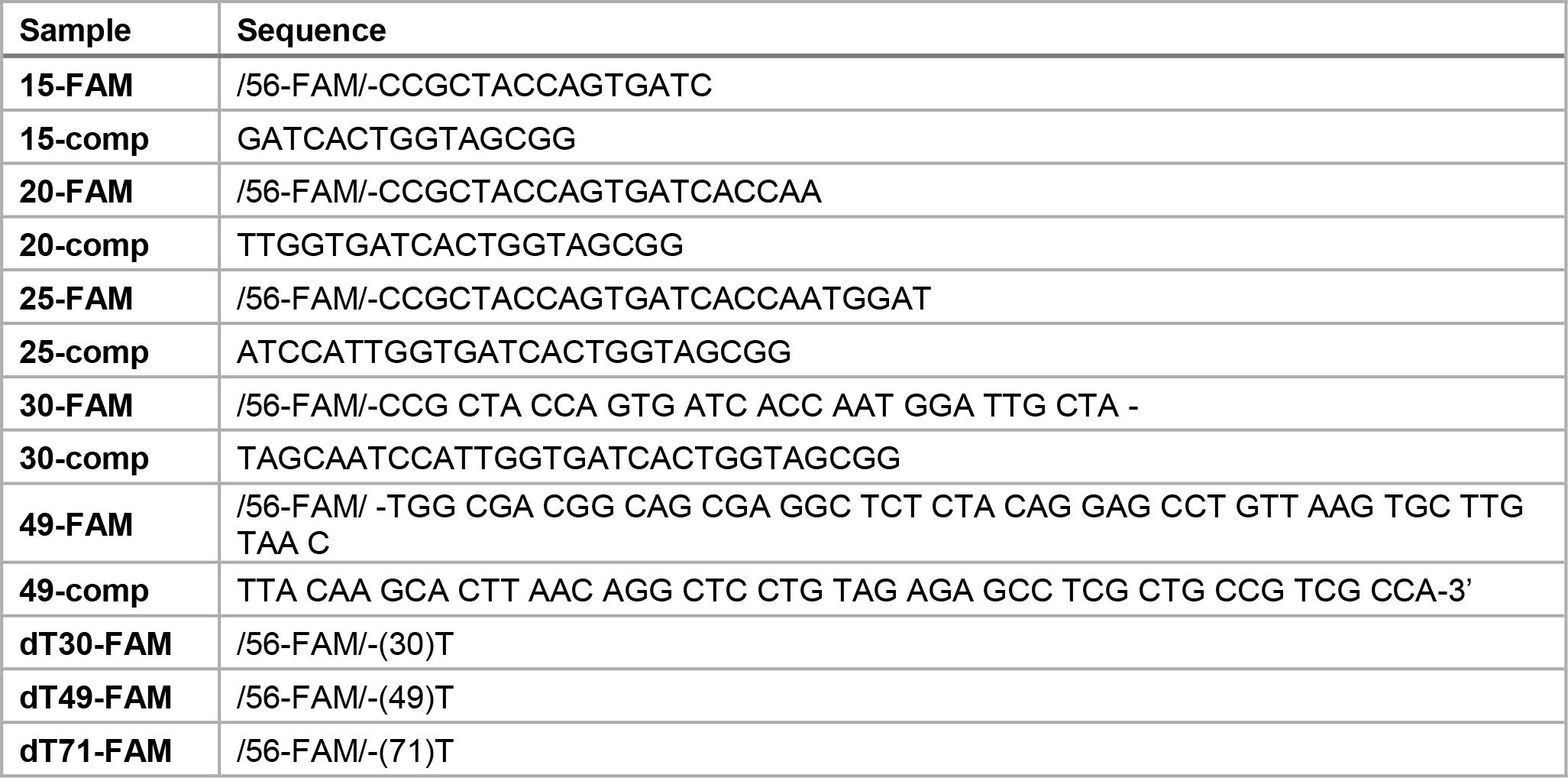
Substrates for DNA binding assay.

**Table S3.**
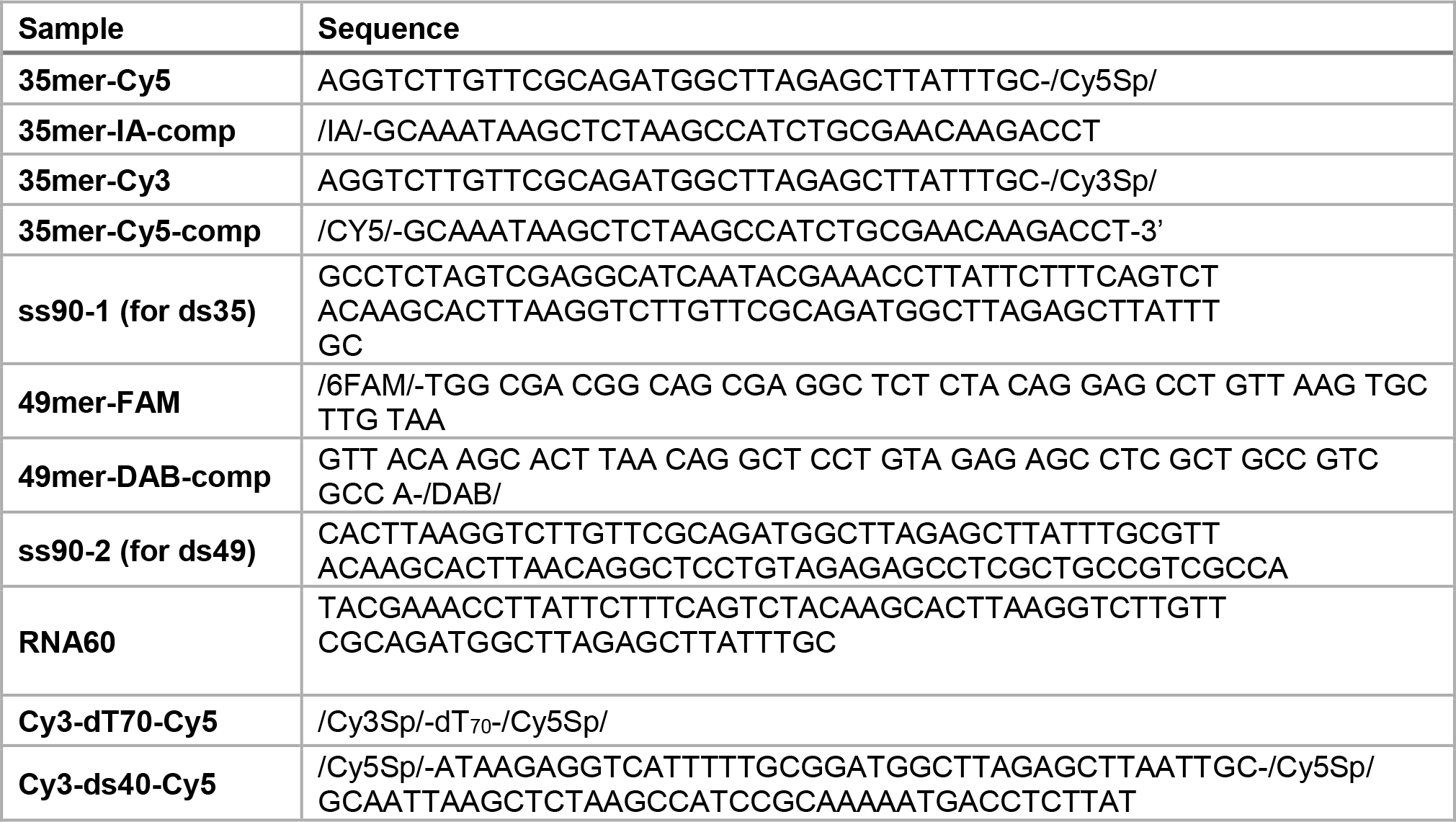
Substrates for strand exchange activity and FRET assays.

**Table S4.**
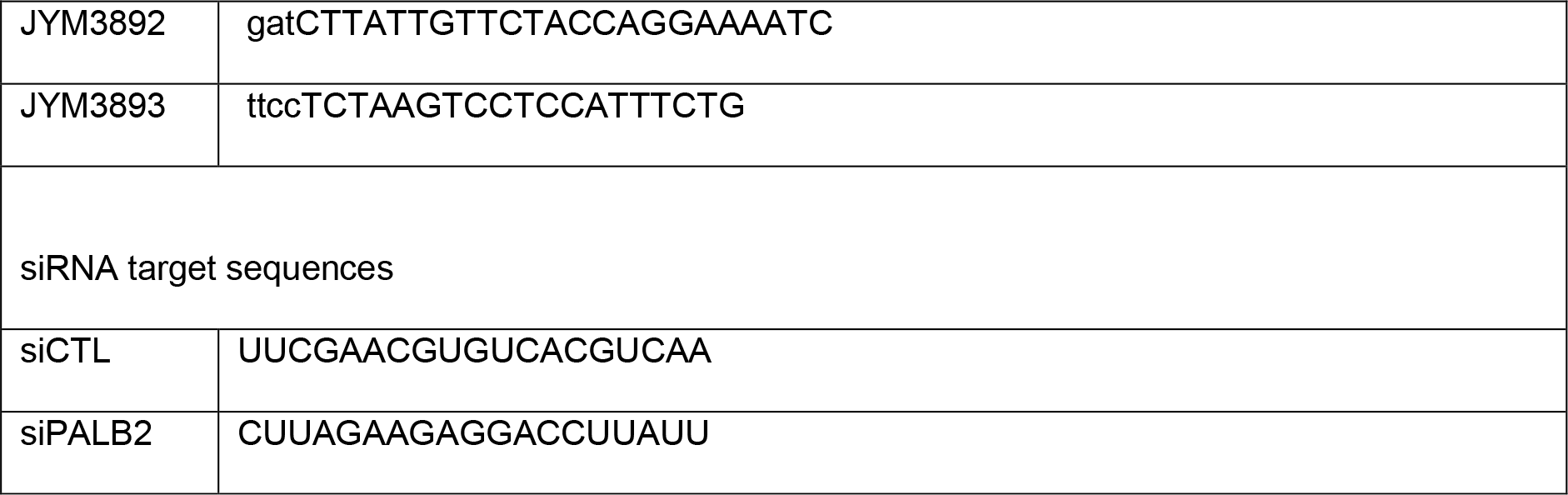
siRNA resistance primers.

**Figure S1.**
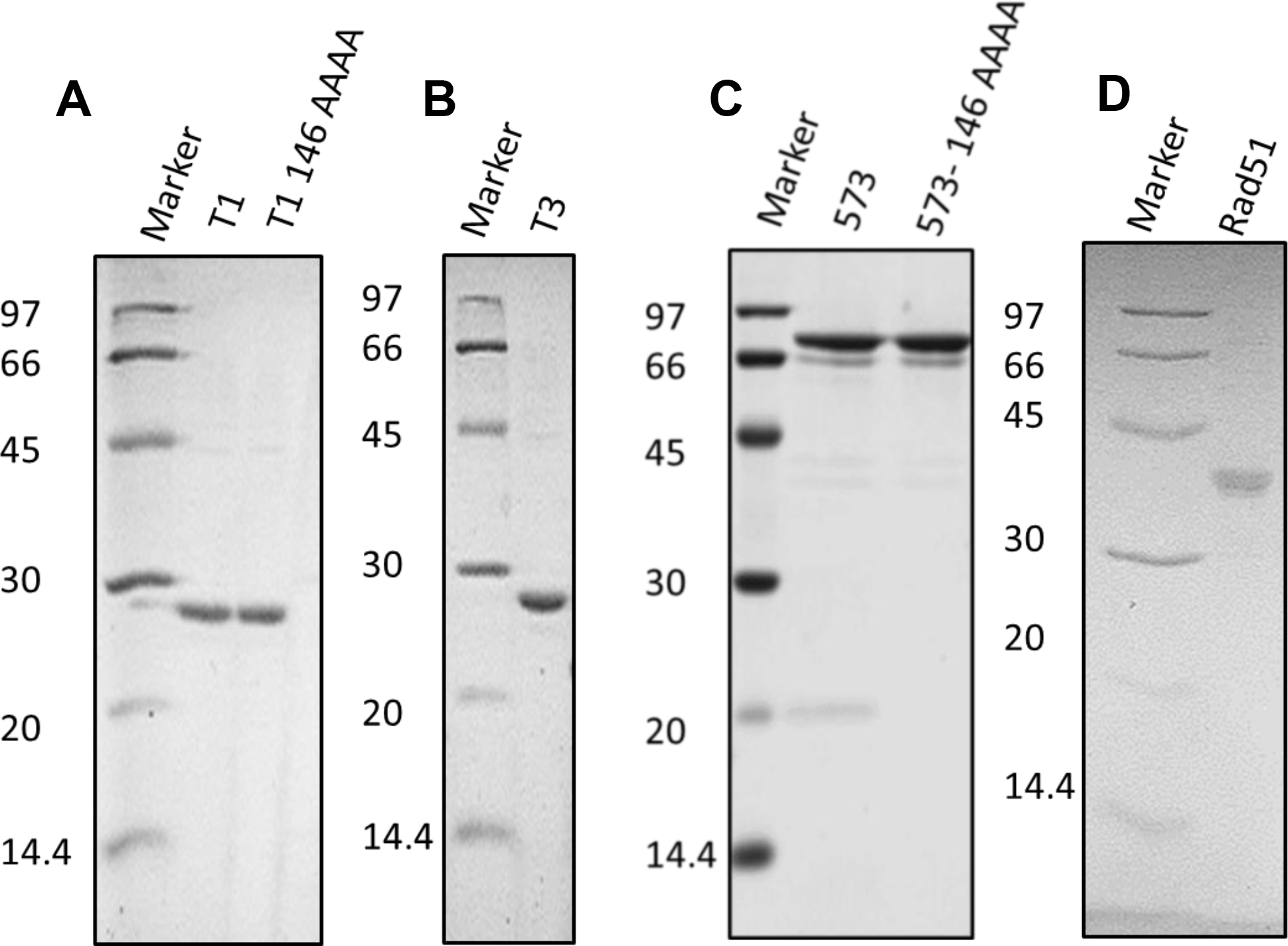
SDS PAGE analysis of purified proteins used in this study: **A)** T1 and T1 146AAAA mutant, **B)** T3, **C)** PB2-573 and PB2-573 146AAAA, and **D)** RAD51.

**Figure S2.**
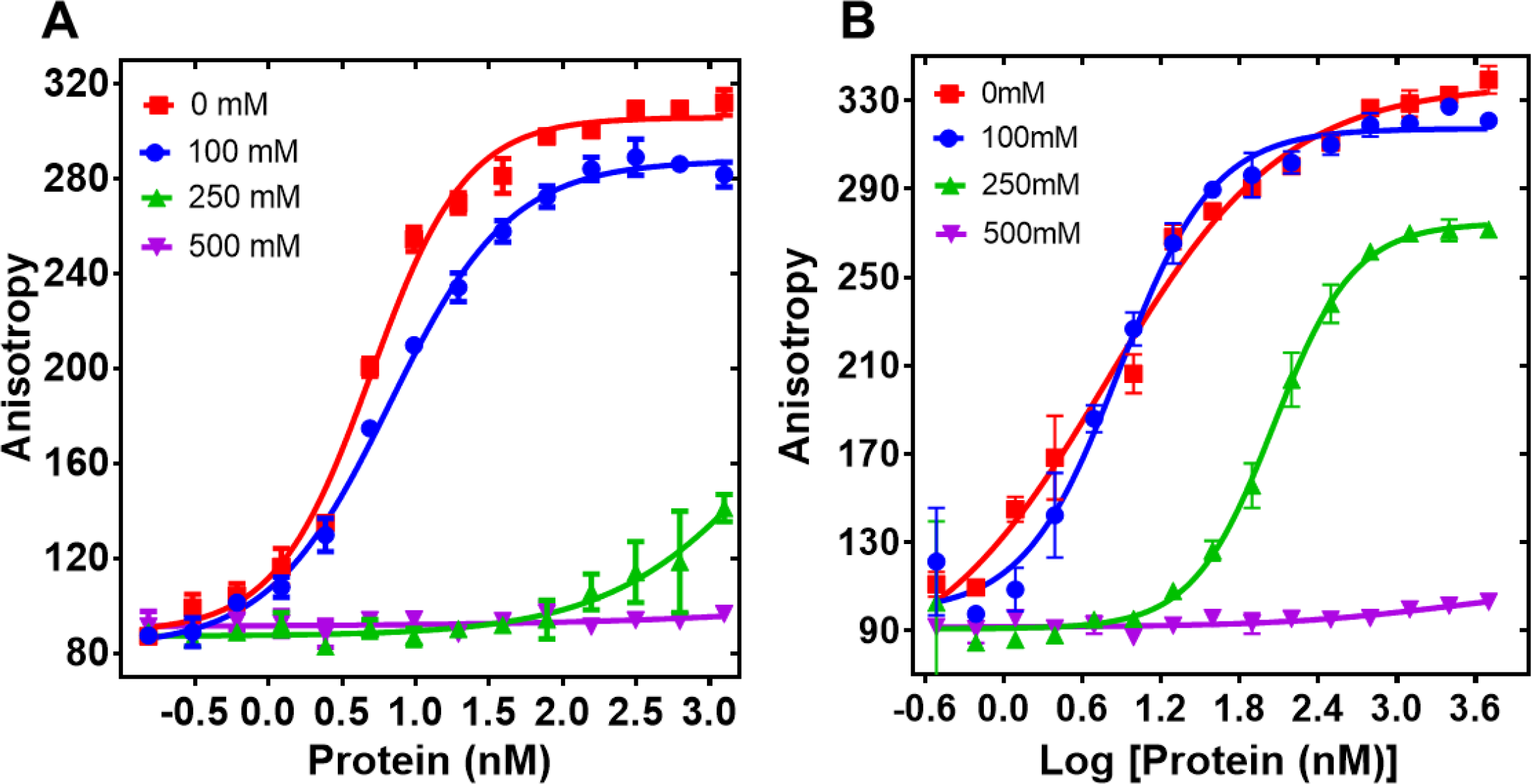
Effect of increasing salt concentration on PALB2 fragments binding to DNA. **A)** Equilibrium binding of T1 with FAM-ss49, **B)** equilibrium binding of PB2-573. Each data point is an average of six readings from two different experiments.

**Figure S3.**
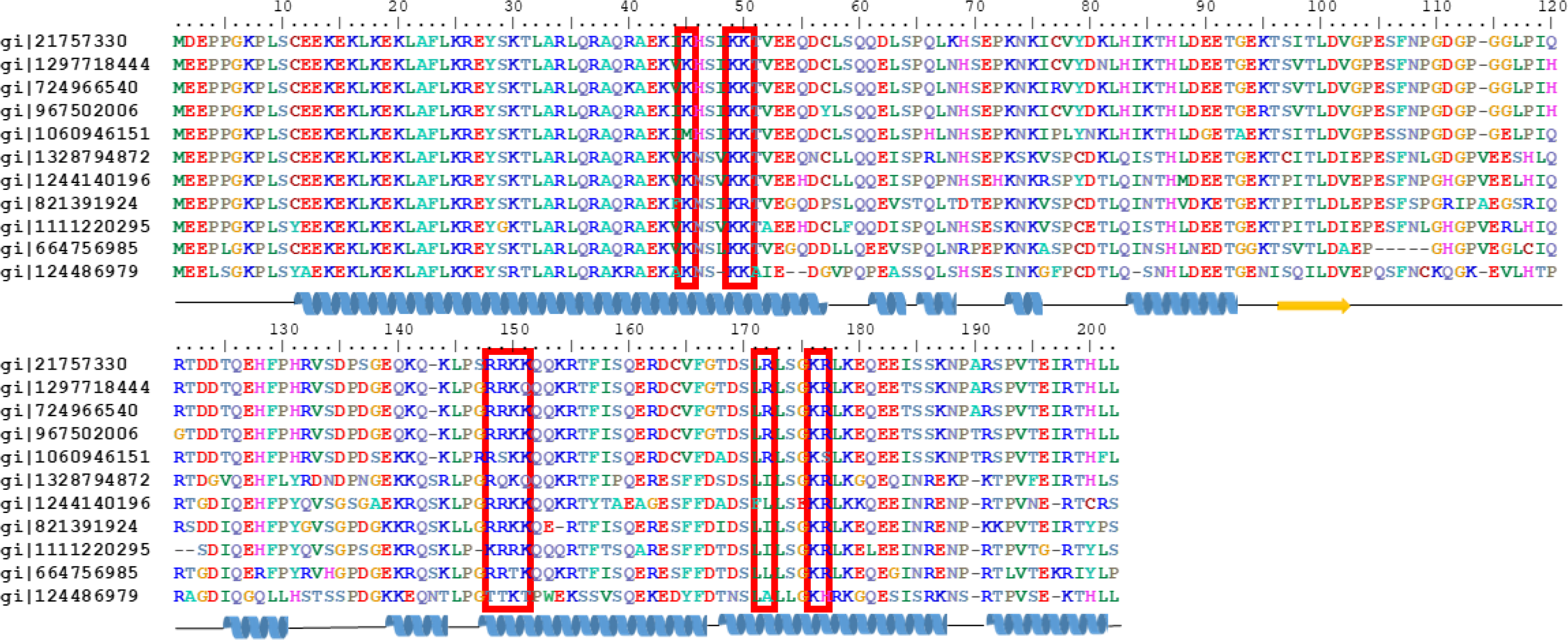
Amino acid sequence alignment of PALB2 T1 from different organisms with residues color-coded accordingly to polarity, with mutated residues identified by red boxes, and with the secondary structure elements depicted at the bottom of alignment in cartoon representation as predicted by Phyre server. Following sequences were used in the analysis. gi|21757330 – Homo sapiens, gi|1297718444 - Piliocolobus tephrosceles, gi|724966540 - Rhinopithecus roxellana, gi|967502006 - Macaca mulatta, gi|1060946151 - Callithrix jacchus, gi|1328794872 - Loxodonta africana, gi| 1244140196 - Enhydra lutris kenyoni, gi|821391924 - Orcinus orca, gi|1111220295 - Panthera pardus, gi|664756985 - Equus przewalskii, gi|124486979 - Mus musculus.

**Figure S4.**
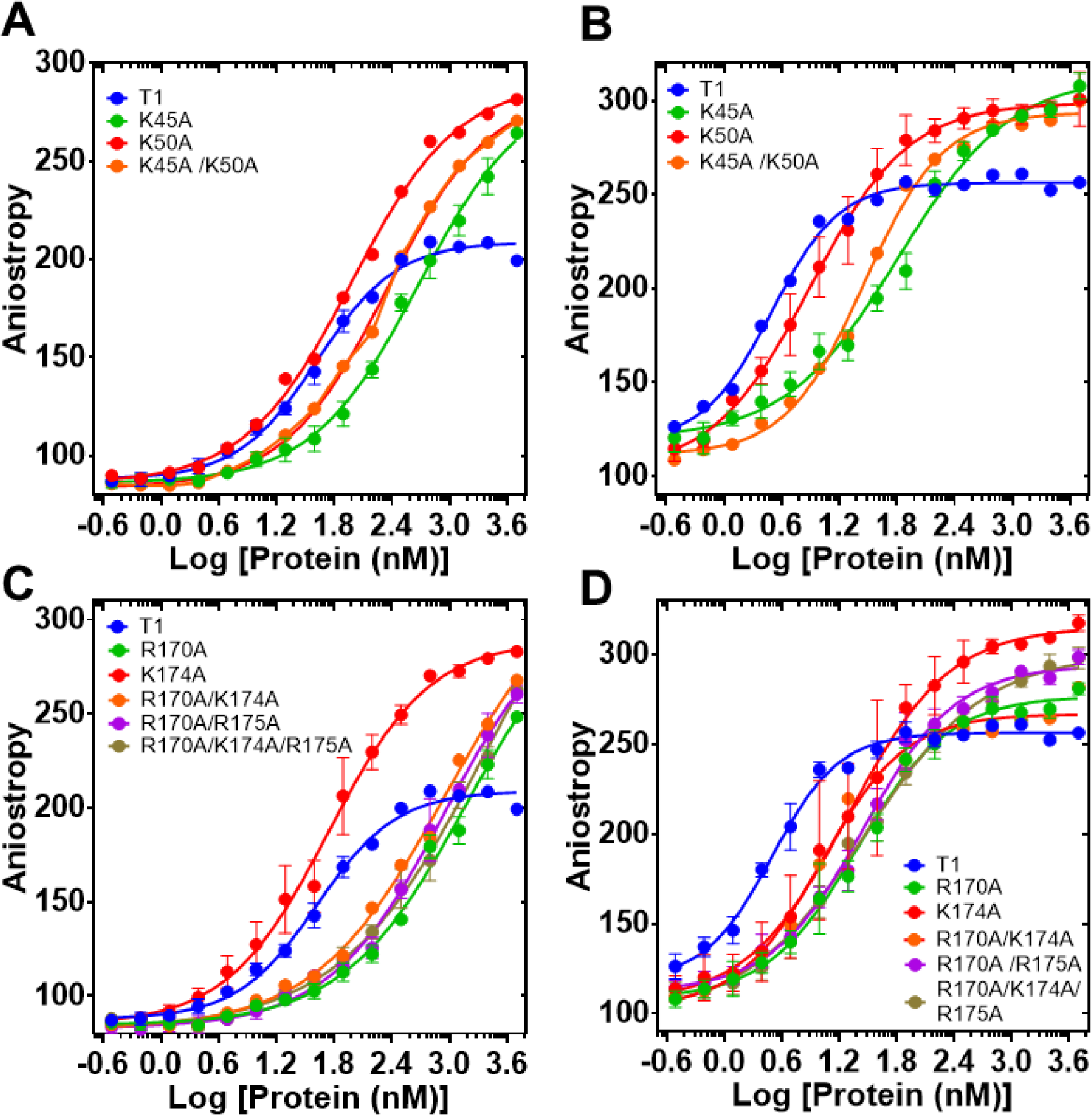
DNA binding of T1 mutants. Equilibrium binding of 5 nm FAM-ss25 (A,C) or FAM-ss49 (B,D) to PALB2 T1 (blue). In A) and B) isotherm for K45A is in green, for K50A in red, for K45A/K50A in black. In C) and D) isotherm for R170A is in red, for K174A in green, for R170A/K174A in magenta, for R170A/R175A in orange, and for R170A/K174A/R175A in black. The binding buffer is 20 mM Tris-acetate pH 7.0, 100 mM NaCl, 5% glycerol, 10% DMSO.

**Figure S5.**
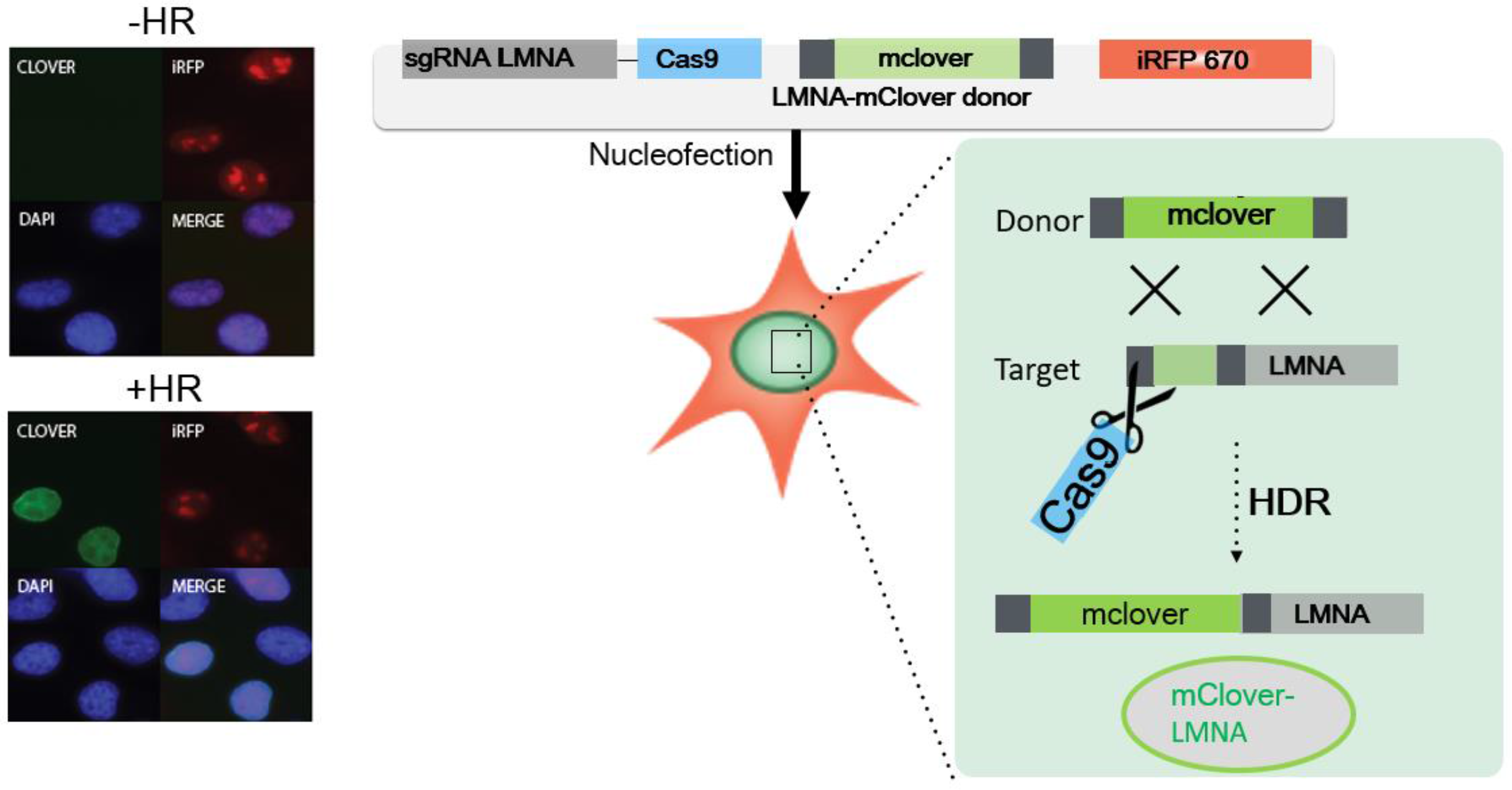
Representative images and **s**chematic representation of CRISPR Cas9/mClover-LMNA1 mediated HR assay. Following nucleofection, Cas9 creates a double-strand break in the LMNA locus leading to integration of the mClover gene by homologous recombination. Clover-labeled Lamin A/C proteins exhibit green fluorescence enriched at the nuclear periphery, which is indicative of the successful gene targeting by homologous recombination.

**Figure S6.**
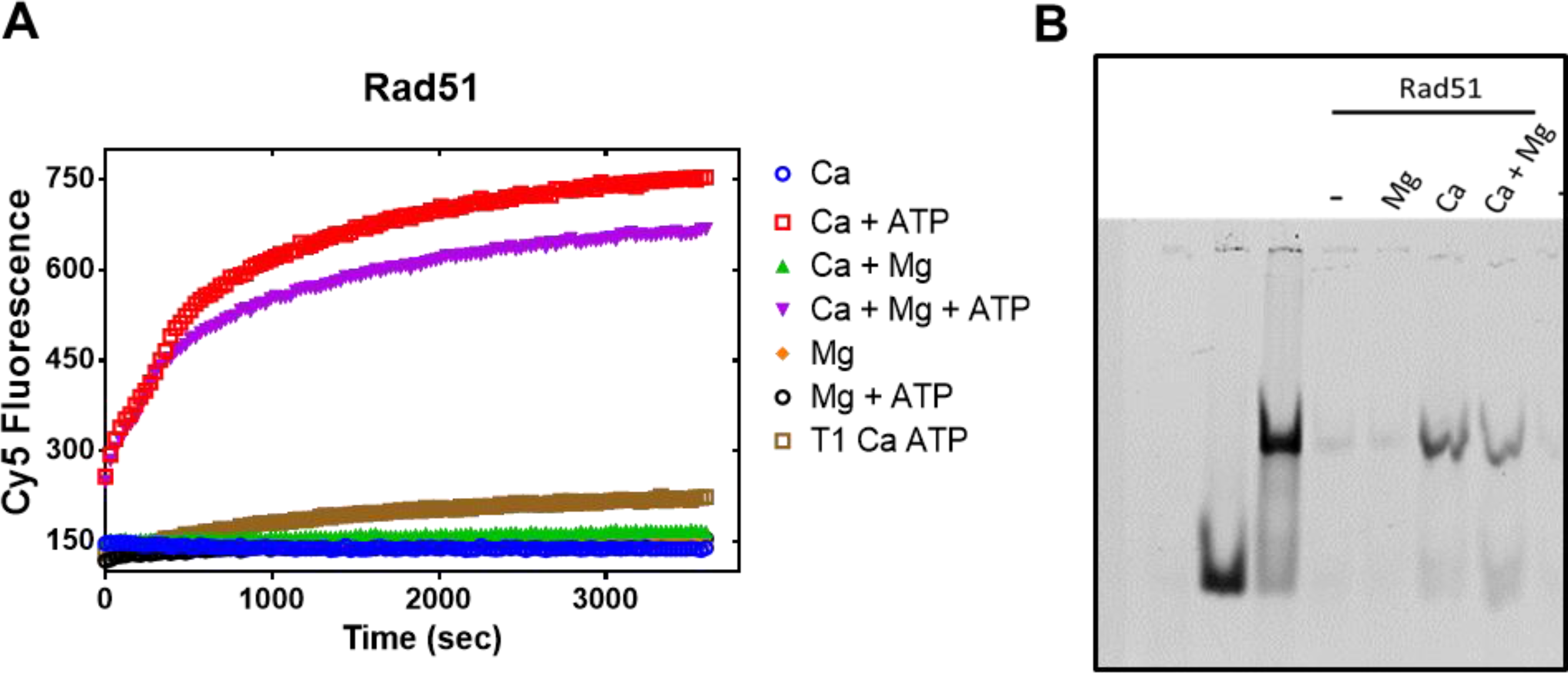
**A)** Strand exchange by RAD51 monitored by Cy5 fluorescence under optimized conditions with 5 mM CaCl_2_, 5 mM MgCl_2_. **B)** End products of the reaction in A) analyzed with EMSA.

**Figure S7.**
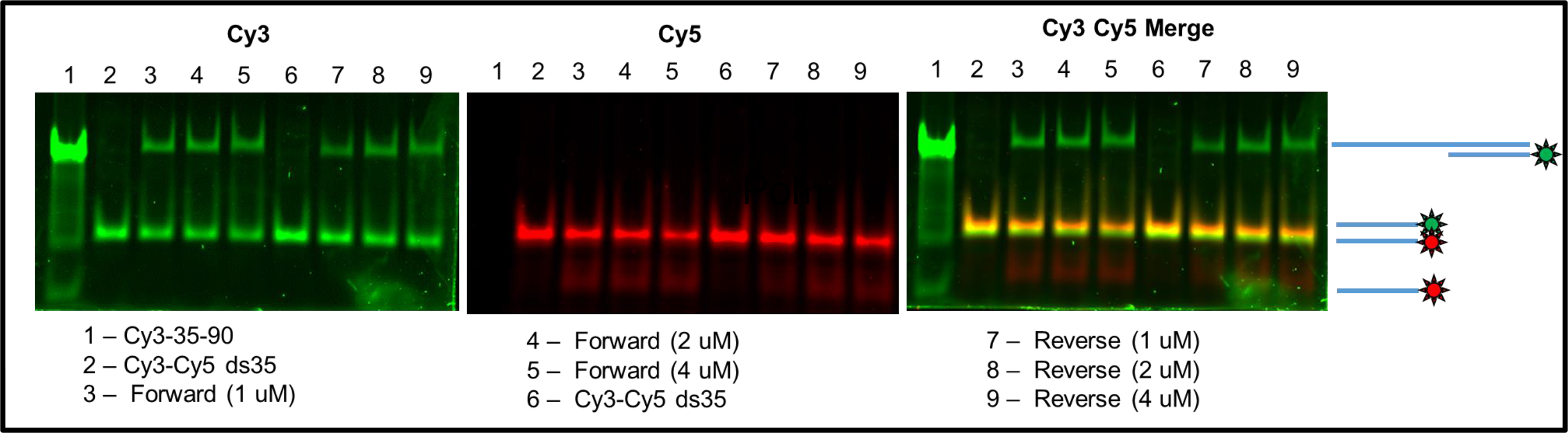
EMSA of N-DBD-mediated strand exchange products using Cy3- and Cy5-labeled ds35 DNA.

**Figure S8.**
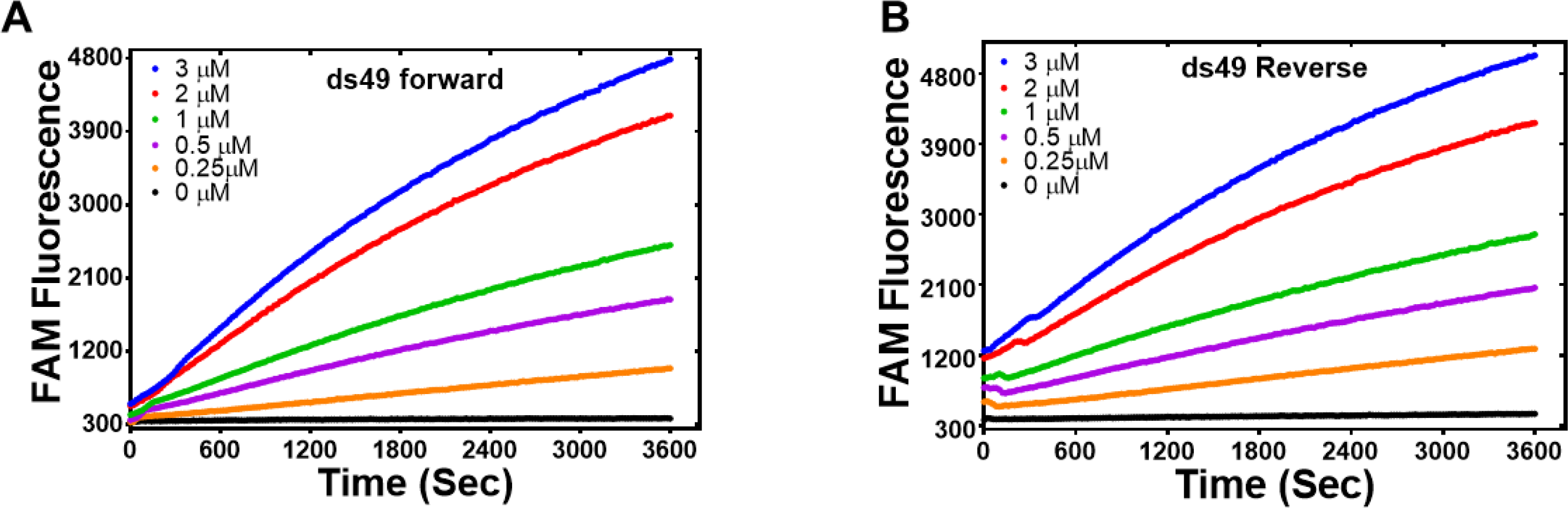
Forward **(A)** and inverse **(B)** strand exchange reactions supported by N-DBD at different concentrations ranging from 0.25 mM to 3 mM performed with ss90 and FAM/Dadsyl-labelled ds49.

**Figure S9.**
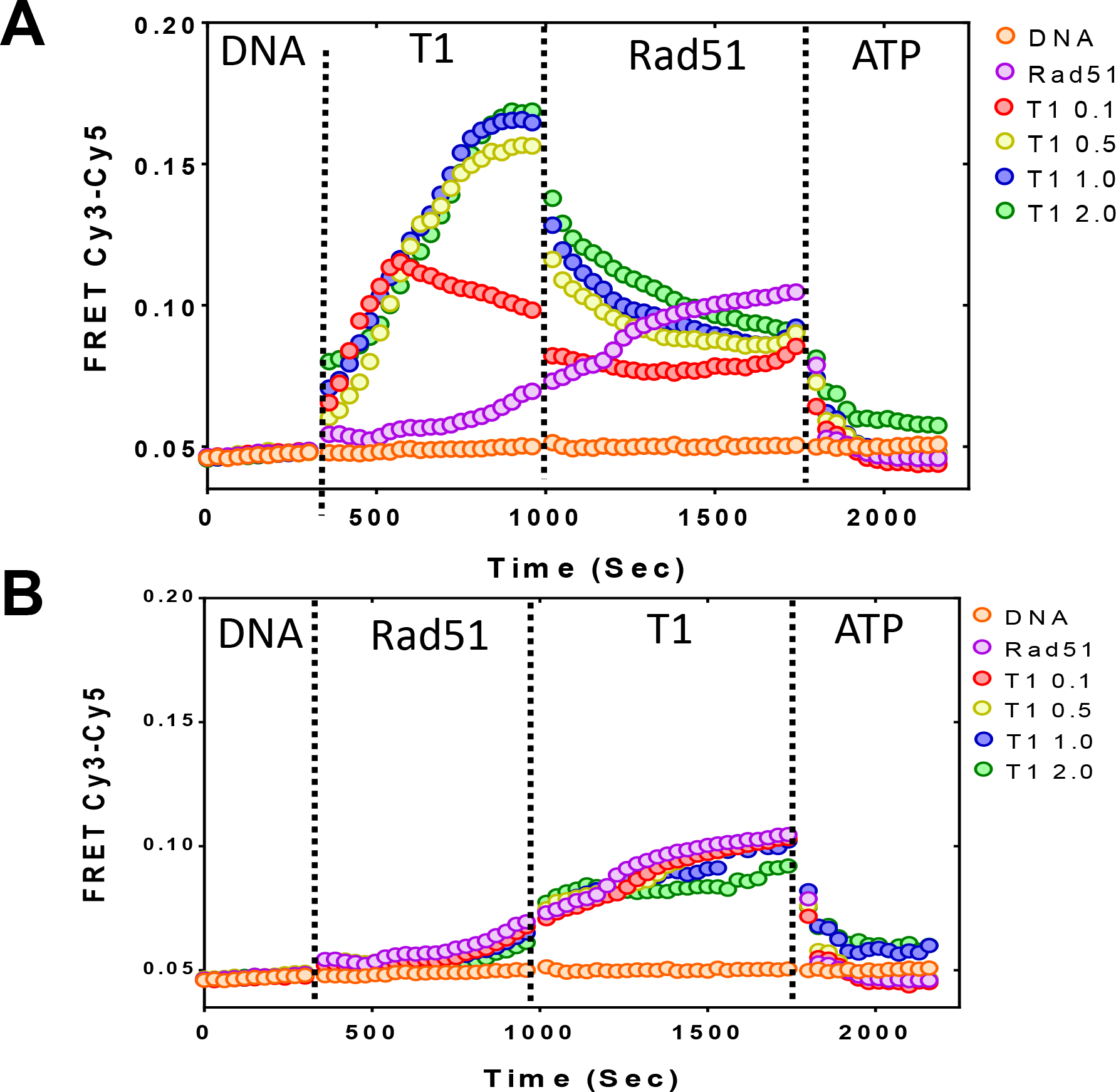
Interaction of PALB2 and RAD51 with ssDNA. **A)** FRET of Cy3-dT_70_-Cy5 alone (initial 5 min) is changed upon addition of different amounts of N-DBD (color-coded) and then by addition of 2 μM of RAD51 in the absence of ATP. FRET decreases further on addition of 2 mM ATP. **B)** A Similar experiment but with RAD51 added first followed by the addition of PALB2 N-DBD.

**Figure S10.**
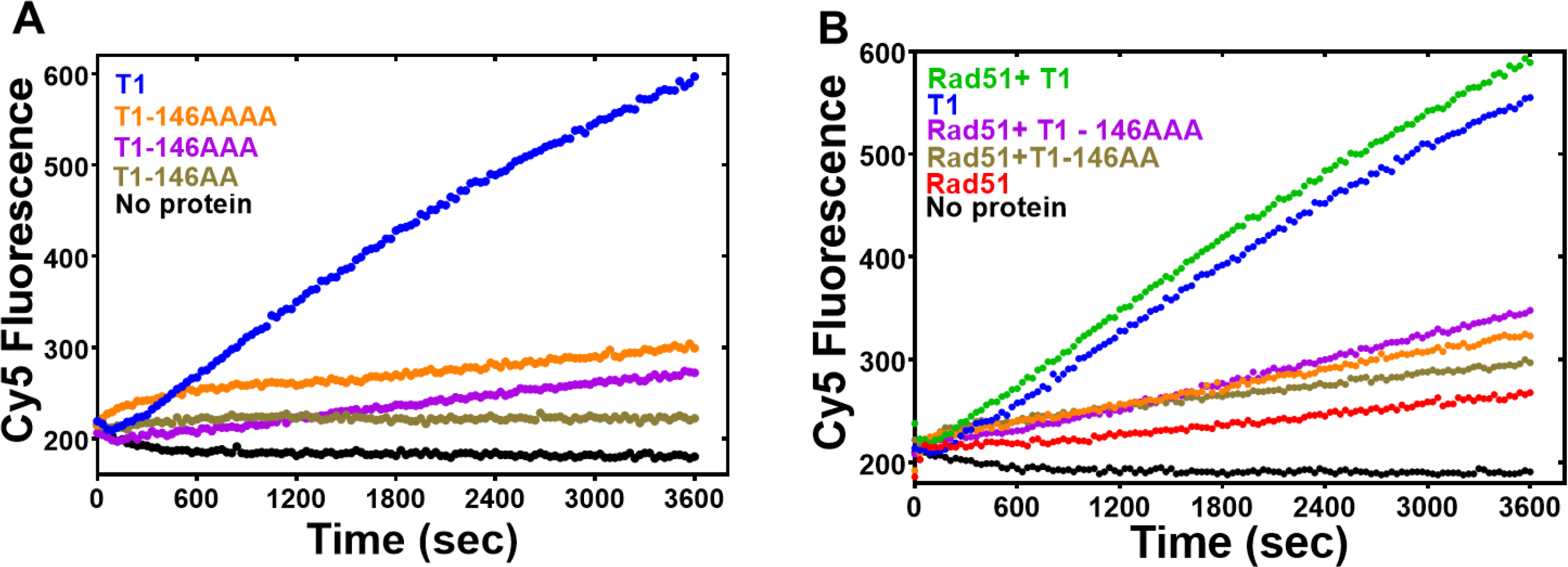
PALB2 DNA binding site mutants do not support strand exchange. **A)** Forward strand exchange activity of PALB2 N-DBD DNA binding mutants. **B)** The strand exchange of PALB2 N-DBD mutants in the presence of Rad51. Strand exchange reactions were performed with Cy5 and Iowa labelled 35bp DNA at 2 μM protein concentration.

**Figure S11.**
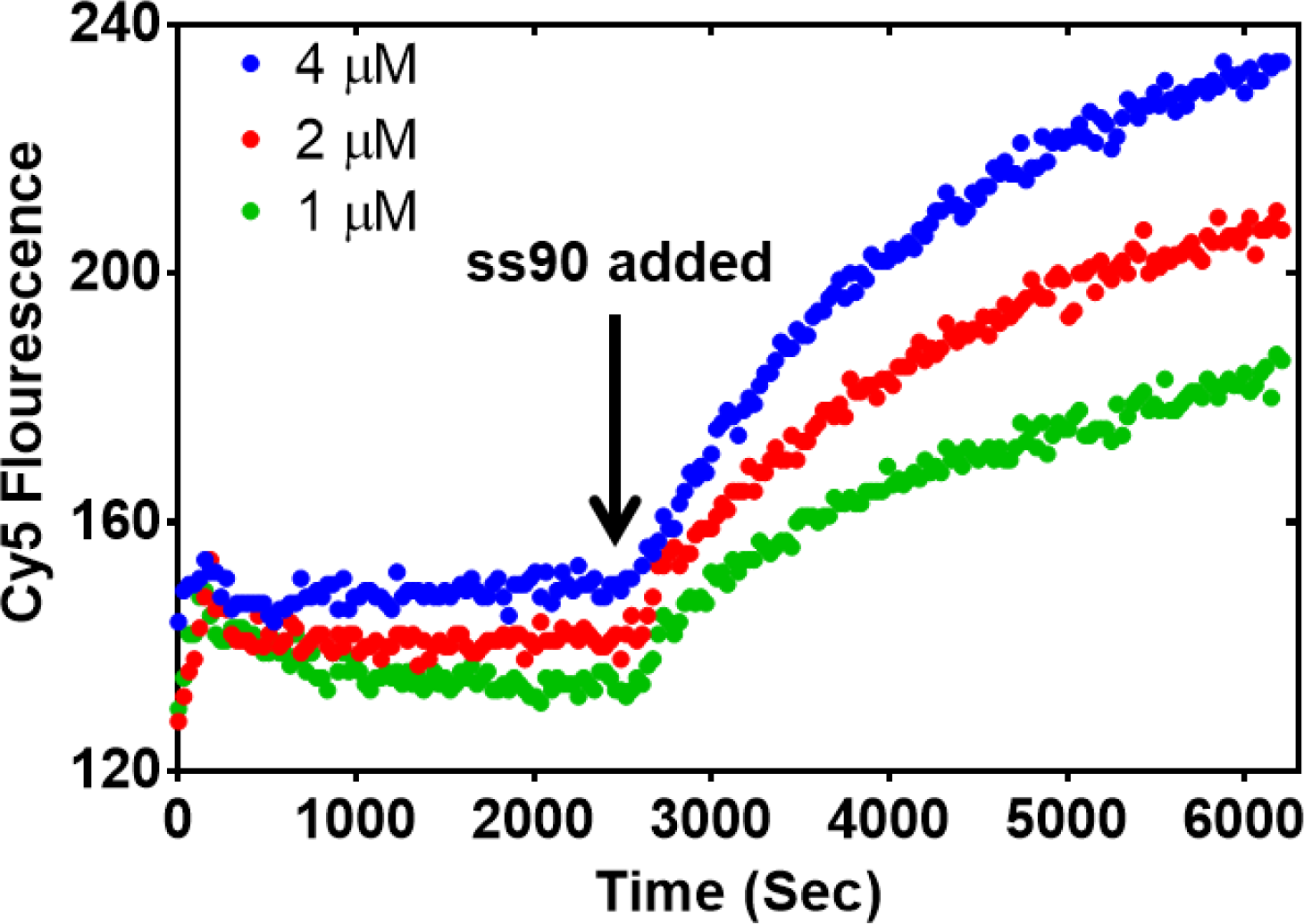
PALB2 does not unwind dsDNA. Cy5/Iowa ds35 DNA was incubated with three different concentrations of N-DBD for 30’ before adding complementary ss90 DNA.

**Figure S12.**
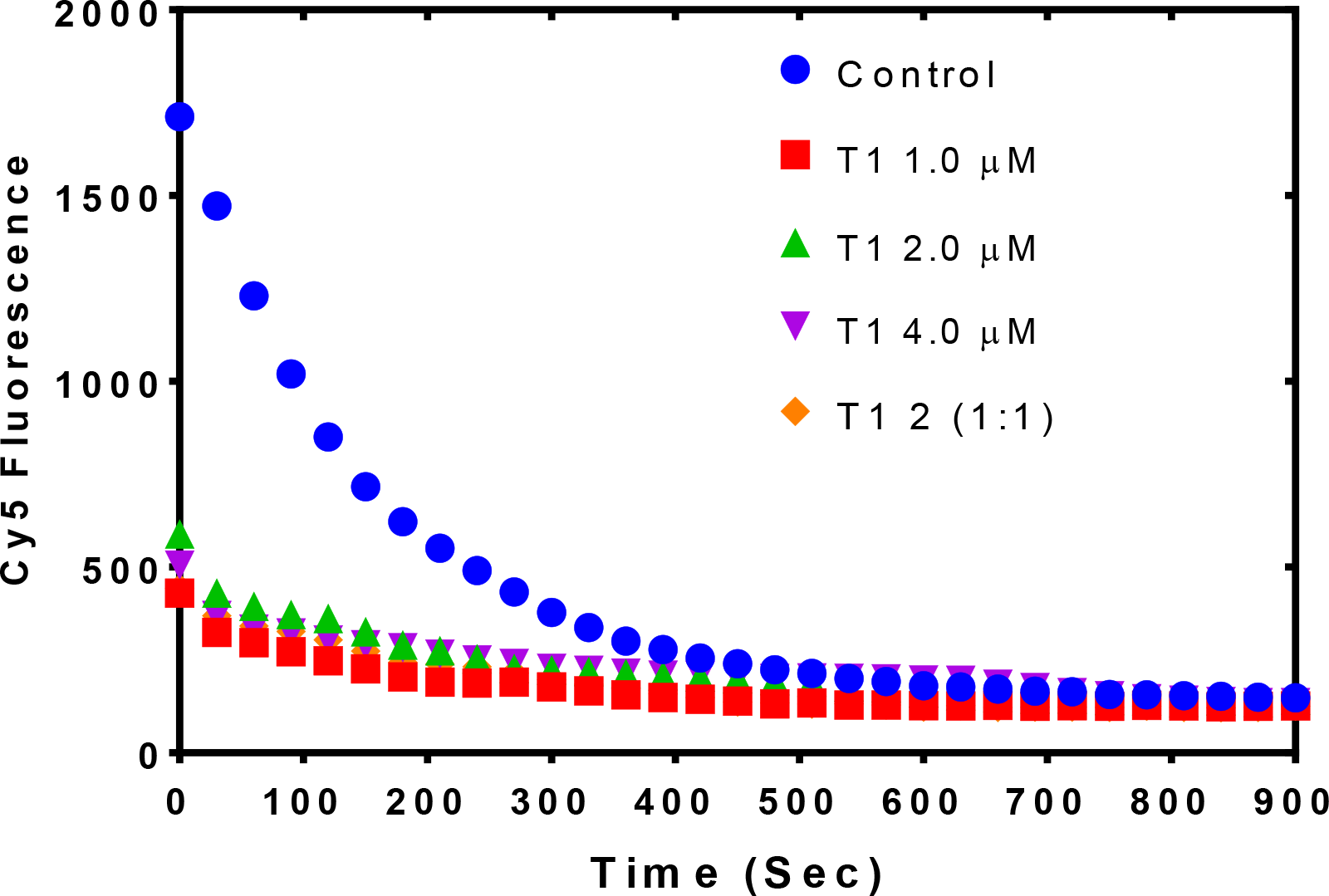
PALB2 anneals complimentary ssDNA. Annealing of Cy5- and Iowa-labeled complimentary ss35 strands in the presence of different concentrations of PALB2 N-DBD.

**Figure S13.**
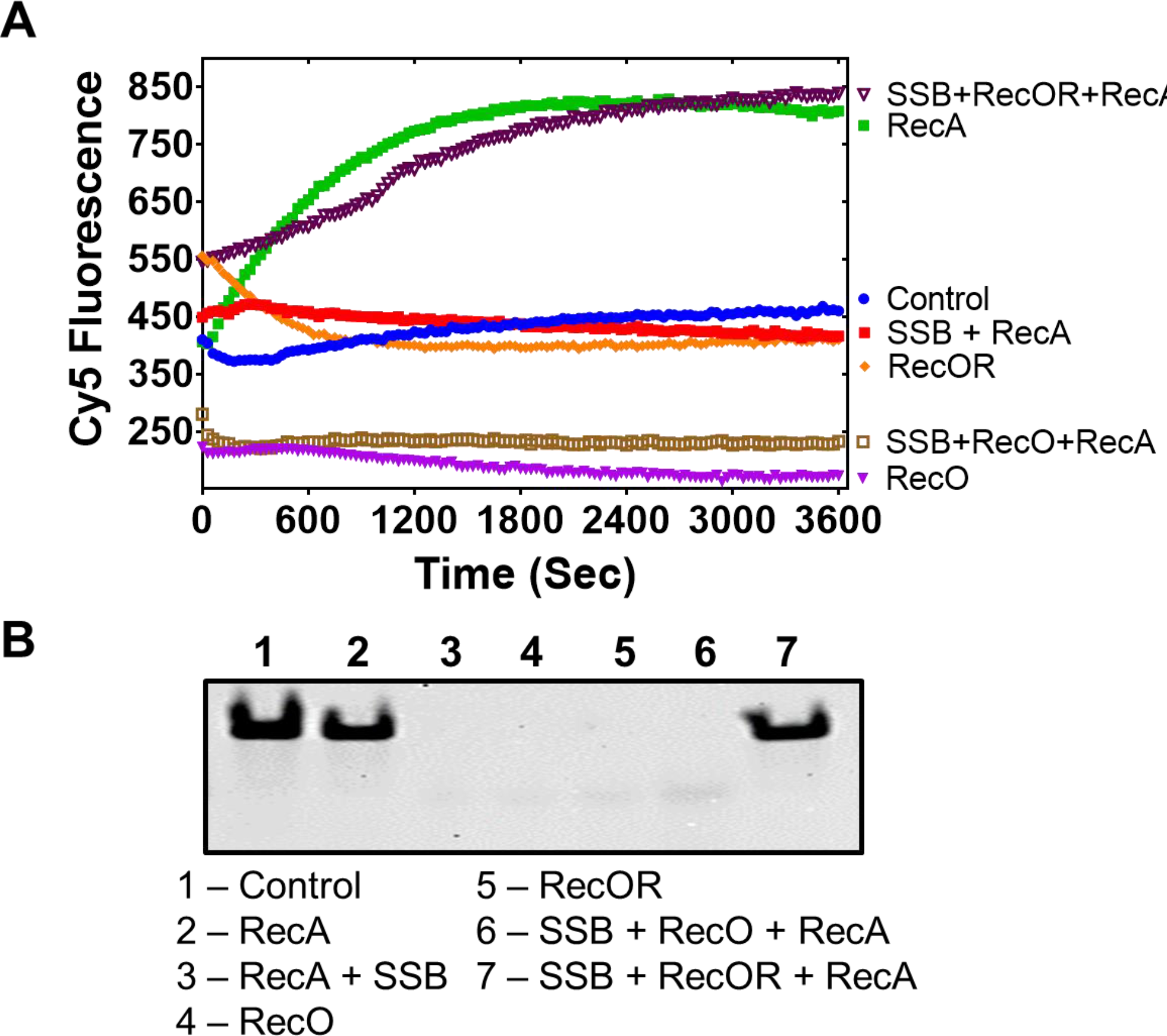
RecO and RecOR do not support strand exchange without RecA. **A)** Strand exchange activity measured under conditions identical to those in Fig. 4 of RecO (magenta), RecOR (orange) and RecOR in the presence of RecA and SSB (brown) monitored by Cy5 fluorescence. Activity of RecA alone without SSB is shown in green. **B)** Reaction products from A) were deproteinized and separated on native PAGE gel.

**Figure S14.**
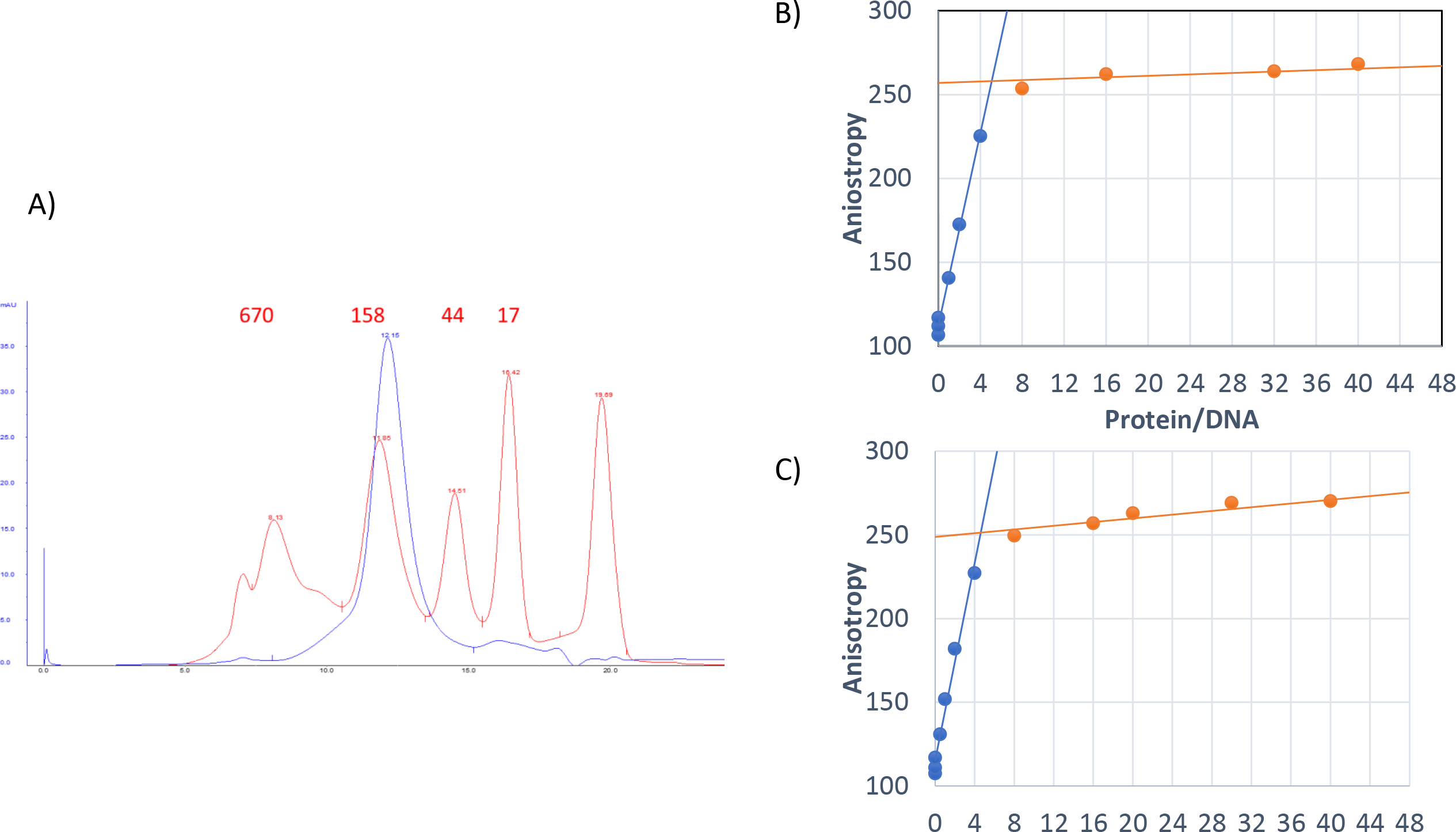
Oligomerization and DNA-binding stoichiometry of PALB2 T1. **A)** Gel filtration of T1 fragment using Superdex-200 10/300 column (blue isotherm). Magenta isotherm corresponds to SEC of BioRad MW standards with MW in kDa shown above each peak. The elution volume of N-DBD correspond to tetrameric or pentameric structure. **B**) and **C**) Titration of FAM-labeled ss49 (at concentration of 50 nM in B) and 100 nM in **C**)) by N-DBD revealed a 5:1 protein:DNA stoichiometry.

**Figure S15.**
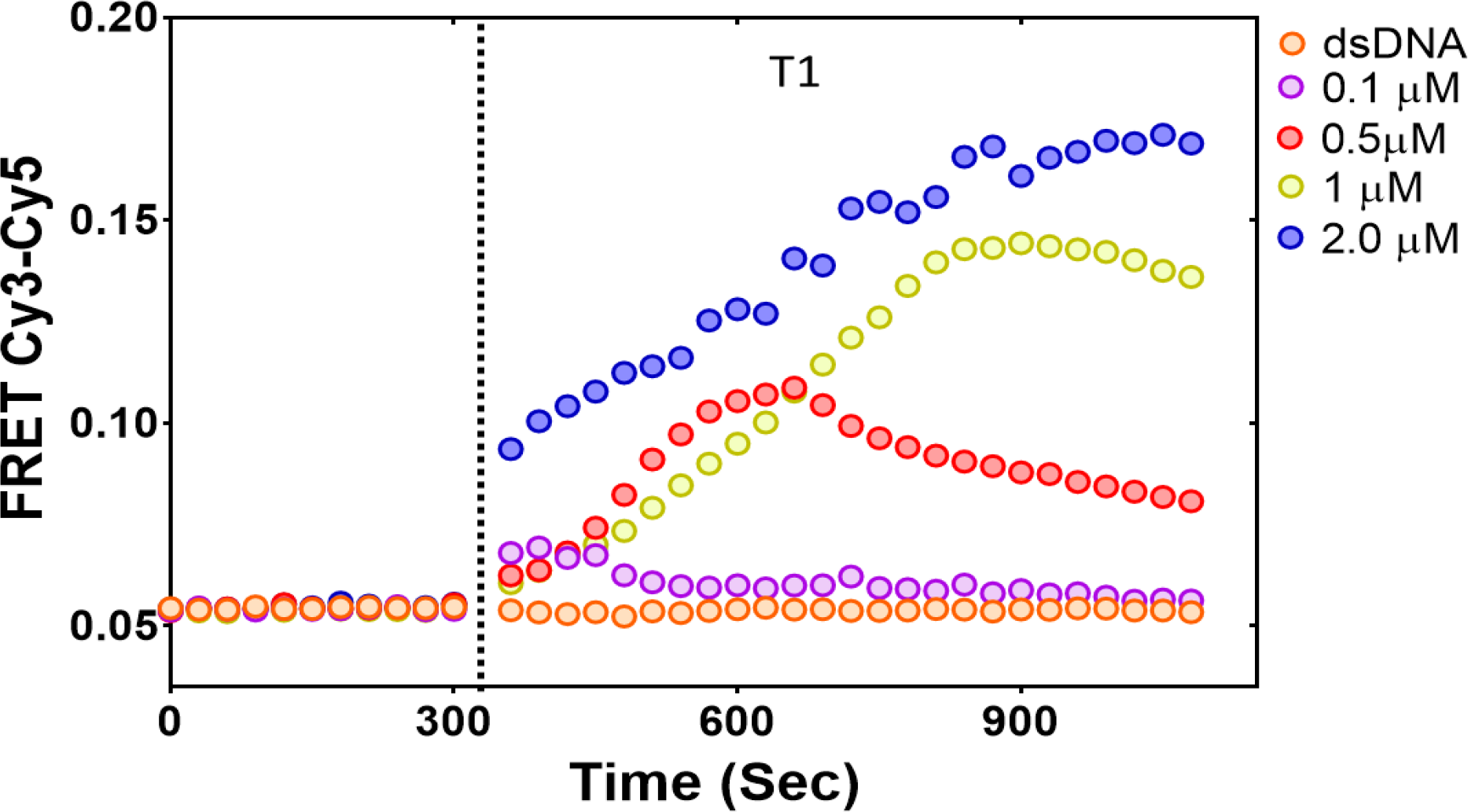
PALB2 N-DBD bends dsDNA. FRET of Cy3-ds40-Cy5 alone (initial 5 min) is changed upon addition of different amounts of N-DBD (color-coded). Protein interaction does not significantly change fluorescence with DNA substrates labelled by either fluorophore alone.

